# 3D Visualization and Proteomic Analysis of Human Cardiac Transthyretin Amyloidosis Tissue Reveals Microangiopathy and Capillary Occlusion

**DOI:** 10.64898/2026.02.09.704431

**Authors:** Joseph P. Donnelly, Jan-Hannes Schäfer, Leonard Yoon, Lynee Massey, Carl Ash, Zi Gao, Karina Nugroho, Marcus Jäger, Zhengyuan Pang, Robert T. O’Neill, Mathew S. Maurer, Evan T. Powers, Gabriel C. Lander, Li Ye, Jeffery W. Kelly

## Abstract

**Background:** Transthyretin amyloidosis (ATTR) is a progressive, degenerative disease affecting the heart and other organ systems, as well as the peripheral, autonomic, and central nervous systems. Although pharmacological and genetic evidence establishes aggregation as a driver of ATTR pathology, the mechanism by which aggregation compromises post-mitotic tissue function is poorly understood. We utilized bottom-up proteomics on wild-type (WT) human cardiac (WT/WT genotype) and V122I human cardiac (V122I/WT genotype) tissue, combined with tissue clearing technology to create an optically transparent tissue architecture to visualize three-dimensional relationships, to better understand TTR cardiomyopathy (CM).

**Methods:** Flash-frozen 0.5 mm cardiac tissue slices from human subjects with end-stage WT-TTR CM, end-stage V122I CM, and slices from an age-matched human control were used for these experiments. Fibril extraction from diseased tissue followed published protocols. Strong denaturant-mediated proteome tissue extraction on samples from each subject facilitated bottom-up proteomics by using liquid chromatography (LC)-mass spectrometry (MS)/MS. Tissue clearing was performed on 0.5 mm cardiac slices utilizing a lauryl sulfate-based lipid removal strategy. Slices were stained using indirect immunofluorescence with antibodies to protein targets identified by proteomics. We used an antibody to non-native TTR and AmyTracker 480 (an oligothiophene dye that binds to amyloid fibrils) to image TTR deposits. ATTR fibrils were characterized structurally using cryogenic electron microscopy (cryo-EM) followed by helical reconstruction.

**Results:** Proteomic cardiac analysis afforded high spectral counts for transthyretin (TTR) and proteins typically associated with amyloid fibrils, e.g. serum amyloid P (APCS). Fibril and cardiac homogenate proteomics revealed high levels of angiogenic and hemostatic proteins, including those composing the complement and coagulation cascades. 3D imaging revealed loss of normal microvascular architecture in CM samples with regions of hyper- and hypovascularization. Microvascular obstruction by capillary thrombosis was also observed in CM. ATTR fibrils adopted the common spearhead fold and were decorated with collagen VI (COLVI), an extracellular matrix component.

**Conclusions:** We hypothesize that ATTR CM is a microangiopathy driven by capillary bed thrombo-inflammation and dysregulated angiogenic revascularization. Phenotypic convergence of WT ATTR CM and V122I ATTR CM was observed via proteomics, 3D imaging, and ex vivo fibril characterization by cryo-EM. We provide evidence of capillary thrombosis in ex vivo ATTR CM tissue. Vasodilation and increased capillary permeability expose components of the vascular basement membrane (VBM) to misfolded TTR. These components are known to promote TTR aggregation and stabilize amyloid fibrils in the extracellular space. Congestion of the VBM prevents appropriate revascularization, reducing cardiac exertional capacity over time, leading to heart failure. Our ATTR CM heart tissue proteomics data shows significant overlap with the proteomic profiles of human AD brain tissues, revealing key amyloid, coagulation, complement, and angiogenesis proteins being changed in amyloidoses.

## Introduction

Transthyretin amyloidosis (ATTR) is a progressive degenerative disease, often initially affecting the heart and/or the peripheral and autonomic nervous systems, and eventually the central nervous system.^1,2^ This disease can be inherited (autosomal dominant) or can occur sporadically as an apparent consequence of aging. When principally affecting the heart, if untreated, ATTR cardiomyopathy (CM) progresses to heart failure with preserved ejection fraction and eventually death in approximately 5-7 years.^2,3^ Pharmacological and genetic evidence support the hypothesis that the post-mitotic tissue degeneration in ATTR is caused by aggregation of the human plasma protein transthyretin (TTR).^2,3^ Kinetic stabilizers such as tafamidis increase the energetic barrier for TTR tetramer dissociation, the rate-limiting step for TTR aggregation, through native state kinetic stabilization, thereby inhibiting aggregation of TTR.^4,5^ Tafamidis dosing thus slows down the progression of both hereditary and sporadic WT ATTR cardiomyopathy by inhibiting aggregation of newly biosynthesized TTR.^4^

While TTR primarily functions as the main carrier of holo-retinol binding protein and as a secondary carrier for thyroxine,^6,7^ studies have also intimated TTR’s ability to regulate angiogenic functions, including capillary growth and endothelial cell proliferation in lung tumors via Akt.^8,9^ The TTR tetramer has been shown to increase expression of LRP1,^10^ a receptor that plays a pivotal role in organized angiogenesis and acts, in part, via Akt.^11^ LRP1 also has a role in the neuronal internalization of amyloidogenic proteins such as tau^12^ and α-synuclein,^13^ as well as participating in TTR-Aβ trafficking,^14^ indicating a potential role for LRP1 in ATTR development. Interestingly, V30M TTR has been shown to downregulate many pro-angiogenic genes in endothelial cell culture,^9^ suggesting TTR tetramer stability may play a role in TTR’s effectiveness in regulating angiogenesis. Indeed, mouse models of Alzheimer’s disease treated with TTR tetramer stabilized with iododiflunisal displayed decreased vascular pathology compared to controls.^10^ Further evidence suggesting a potential role of the native tetrameric structure in TTR’s functional angiogenic properties is seen in the gradual increase in circulating endothelial progenitor cells in cardiac ATTR patients treated with the TTR stabilizer tafamidis.^15^

The pathophysiology of ATTR CM remains unclear. One hypothesis is that TTR amyloid fibril accumulation in the myocardium leads to a restrictive or infiltrative CM, wherein muscle tissue in the heart’s lower chambers (ventricles) becomes stiff, and the ventricles cannot fill with blood efficiently, leading to reduced flow of blood in the heart.^1,16^ Histologically, ATTR amyloid deposition has been described as patchy, i.e. without a discernable deposition pattern, in contrast to the perivascular pattern observed in other forms of cardiac amyloidosis, such as in immunoglobulin light chain amyloidosis (AL).^1,3^ A 2016 histopathological study of both ATTR and AL cardiac amyloidoses described the most common ATTR deposition pattern as discrete pericellular, though nodular and interstitial deposition patterns were also observed.^17^ While vascular deposition was much more prevalent in AL pathology, the study showed ATTR deposition in the intramyocardial vasculature, especially in intramyocardial arteries.^17^

Histopathological ATTR and AL examinations have historically been done by two-dimensional visualization. Clearing methodology, which preserves tissue architecture while removing lipids to render the tissue optically transparent, allows much thicker tissue samples to be imaged gaining 3D insights into pathology.^18^ To our knowledge, this study is the first time that tissue clearing technology has been used to analyze human ATTR CM pathology in an ex vivo 0.5 mm human LV free wall tissue slice.^18^ Previous histological examinations were limited by two-dimensionality, preventing the intricate visualization of small structures such as capillary vessels. Unlike cryo-electron microscopy examination of isolated amyloid fibrils from tissue, requiring fibril isolation techniques, cryo-electron tomography has the potential to visualize tissue architecture in its relationship to amyloid fibrils; however, this approach is technically demanding.^19^ Tissue clearing technology, employing a uniform refractive index across the sample allowing for z-stack imaging of dense tissue and 3D reconstruction of physiological structures of interest using fluorescent labeling of strategic proteins, is easier to implement and was used here.

We also employed a bottom-up un-enriched proteomic analysis of explanted ventricular slices, wherein the proteome was extracted under strongly denaturing conditions overnight utilizing 6M guanidine thiocyanate. Proteomics was performed on ventricular slices from two patients, one WT ATTR CM and one heterozygotic V122I ATTR CM, both afflicted with end-stage ATTR cardiac amyloidosis. These proteomic results were compared to age-matched control human ventricular slices, revealing evidence for thrombo-inflammation, with notable elevations in proteins found in the extrinsic coagulation and the classical complement cascades. Proteomic evidence for ongoing angiogenesis and proteins involved in hemostasis were also upregulated in ATTR CM. Tissue clearing of ventricular slices from the respective hearts using sodium lauryl sulfate revealed loss of normal microvascular architecture in ATTR CM, with regions of hypovascularization associated with amyloid deposition, suggesting that amyloidogenesis is associated with a loss of vascularization. Our results provide evidence of capillary occlusion in ex vivo cardiac ATTR tissue. This conclusion is based on the observation of thrombin in vessels measuring < 10 µm in diameter, with loss of vascular architecture downstream of the clot indicating capillary rarefaction. Microvascular occlusion was observed at the periphery of amyloid deposits, which appear to begin in the vascular basement membrane before becoming more diffuse. Attempted revascularization of the occluded vessels, required to restore full functionality to the underlying cardiomyocytes, appeared dysregulated and obstructed by proximal amyloid deposition in the vascular basement membrane/extracellular space. We hypothesize that vasodilation and increased capillary permeability expose components of the vascular basement membrane to misfolded TTR in the extracellular space. Such components are established to nucleate TTR amyloid fibril formation, exacerbating the pathology. We present evidence for such interactions in the form of a low resolution cryo-EM structure of such a component (collagen VI) bound to an ATTR fibril.

## Materials and Methods

### Data Availability

Antibodies used in tissue staining and western blots are listed in Table S1. All density maps and models reported in this article were deposited in the Electron Microscopy Data Bank (EMDB), the Protein Data Bank (PDB), and the Electron Microscopy Public Image Archive (EMPIAR). WT ATTR apex 1 (PDB-9PX6; EMD-71953; EMPIAR-12909), WT ATTR apex 2 (PDB-9PX7; EMD-71960; EMPIAR-12911), V122I ATTR apex (PDB-9PX9; EMD-71962; EMPIAR-12912), Human Collagen VI microfibril (EMDB-73313) and ATTR-Collagen VI Co-fibrils (EMDB-73312). Cryo-EM data collection, refinement and modeling statistics are reported in Table S2. Raw proteomics data can be found in Tables S3, S4, and S5.

### Tissue Samples

Two explanted hearts from genetically tested ATTR CM patients receiving orthotopic heart transplantation for end-stage cardiac ATTR amyloidosis at Columbia University Medical Center were dissected ex vivo, flash frozen in liquid nitrogen, and shipped to Scripps Research Institute on dry ice, where they were stored at -80°C for several years until used for this study. The wild-type ATTR sample contained two large pieces: full width cardiac apex, full width left ventricular (LV) free wall. The V122I ATTR samples contained five separate dissections: LV cardiac apex, LV free wall, interventricular septum, right ventricular (RV) cardiac apex, RV free wall.

The control sample was obtained through the Human Tissue Biorepository at Accio Biobank Online (https://biobankonline.com/). Upon requesting a non-diseased human heart specimen, we were added to an on-call list as the first called after a donor heart was turned down by the United Network of Organ Sharing (UNOS). Inclusion criteria were >55 years old, no known cardiovascular history, and non-cardiac death. The heart we accepted had a warm ischemia time of 80 min. The heart was shipped on ice, and it was dissected and flash-frozen in liquid nitrogen immediately upon reception. The dissected tissue was stored at -80°C until used.

0.5 mm slices of full width ventricular samples from each subject (WT, V122I, control) were used for imaging via tissue clearing.

### Tissue Clearing

Antibodies used in tissue staining and western blots are listed in Table S1. Pilot clearing studies and initial optimization were performed using the hydrogel-reinforced DISCO (HYBRiD) method as described in Nudell et al.^18^ Later studies, including those comprising the data in this manuscript, were performed using Tocris Tissue Clearing Pro reagents following the protocol outlined in Biotechne’s Tissue Clearing Pro Guidebook V1.1, available at https://resources.tocris.com/pdfs/guidelines/tissue-clearing-pro-guidebook.pdf.

Briefly, block samples of full thickness LV free wall were fixed in 4% paraformaldehyde (PFA) solution. Fixed samples were sliced into 0.5 mm full thickness sections using a vibratome and stored in PBS with 0.02% sodium azide until ready for tissue clearing. The tissue clearing protocol began with two PBS washes, followed by overnight incubation at room temperature (≈ 25 °C) using the Tissue Clearing Pro Permeabilization Buffer (6% Sodium lauryl sulfate; Tocris Bioscience), recommended for dense tissue. The sample slices were again washed in PBS twice before being subjected to stepwise dehydration at 4 °C with gentle shaking from 50% ethanol/ PBS, 80% ethanol/DI water, and finally 100% ethanol. Sample slices were then incubated in 20% DMSO/ethanol before rehydration in reverse order. Sample slices underwent incubation in Tissue Clearing Pro Penetration Buffer at room temperature (≈ 25 °C) before being incubated with Tissue Clearing Pro Blocking Buffer at 37 °C with shaking. Slices were then transferred to the primary antibody solution (1:500 dilution per antibody [Table S1] in PBS with 0.1% Triton X100) at 37 °C with shaking. Following primary antibody incubation, samples were washed 10 times in PBS with 0.1% Triton X100 and transferred to the secondary antibody solution (1:500 dilution per antibody, 1:1000 dilution AmyTracker480 [Table S1]). Slices were washed 10 times with PBS + 0.1% Triton X100 at room temperature (≈ 25 °C) before undergoing stepwise dehydration as described previously at 4 °C. Slices were then rehydrated with refractive index matching solution Tissue Clearing Pro Reagent 1, followed by incubation with Tissue Clearing Pro Reagent 2 at 4 °C. Slices were mounted on microscope slides using a 500 µm iSpacer and Tissue Clearing Pro Reagent 2.

### Microscopic Imaging and Analysis

Imaging was performed using a Zeiss Celldiscoverer CD7 with a LSM 900 Confocal microscope. Image processing was performed using Arivis Pro software associated with the Zeiss Celldiscoverer CD7.

### Fibril Extraction

Amyloid fibril extraction was performed as described by Annamalai et al.^20^ Briefly, 125-250 mg cardiac tissue (WT-ATTR: LV Free Wall x2, Apex; V122I-ATTR: LV Apex, Interventricular septum, RV free wall) was diced and washed three times in 0.5 mL Tris calcium buffer. Each sample was then incubated overnight at 37 °C in 1 mL of 5 mg/mL Clostridium histolyticum collagenase in tris calcium buffer. The samples were centrifuged at 3100 g for 30 min at 4 °C and the supernatant was discarded. The samples were homogenized in 0.5 mL Tris EDTA buffer, centrifuged at 3100 g for 5 min at 4 °C, and the supernatant was discarded. After 10 cycles of the homogenization step, it was repeated with 0.5 mL ice cold water, with the supernatant stored as fibril extract. Extracts were worked up for mass spectrometry as described, starting with the BCA Protein Assay (Thermo Scientific # 23225) described below.

### Mass Spectrometry Proteomic Analysis

200 mg of ventricular tissue was removed from the three aforementioned heart samples from three separate locations: cardiac apex, LV free wall, and interventricular septum (wild-type ATTR sample used opposing ends of LV free wall along with the cardiac apex). The samples were homogenized in PBS using an Ika-Werk Ultra-Turrax tissue homogenizer. Each sample was then split, with half of the samples incubated overnight in 6M guanidine thiocyanate and the other half undergoing the standard proteome extraction digestion protocol described below.

Protein concentrations of the samples were normalized using Pierce BCA Protein Assay Kit according to the manufacturer’s instructions (Thermo Scientific # 23225). One hundred µg of each sample was transferred in 200 µL urea buffer from Abcam FASP Protein Digestion Kit (Abcam #ab270519) or 200 µL 6M guanidine thiocyanate (Sigma #G9277) onto 3 kDa Microcon filter (Millipore) on the provided collection tube. The guanidine samples were placed at 4 °C overnight. The kit’s Proteome Extract Digestion protocol (Abcam #ab270519) was followed, starting at step 6.3. The 50 µL 200 mM TCEP membrane incubation prior to step 6.4 was omitted. Step 6.4 was modified to add 40 µL of Urea Sample Solution (instead of 90 µL). Step 6.9 was performed with Trypsin Platinum (Promega), 1 µg in Digestion Solution from Abcam FASP kit was added on top of membrane overnight at 37 °C. After step 6.14, digested peptides were acidified with 10 µL of formic acid and dried via speed vac. The peptides were then resuspended in 100 µL of 100 mM HEPES, pH 8.5.

Pierce Quantitative Colorimetric Peptide Assay (Thermo Scientific #23275) was performed according to the manufacturer’s instructions. After quantification, 25 µg of peptide from each sample was labeled with 250 µg of TMTpro 16plex (Thermo Scientific #A44521) in > 40% acetonitrile for 60 min. After labeling, TMT labeling was quenched by 5% hydroxylamine for 15 min. Samples were then pooled and dried via speed vac. Dry samples were resuspended in 300 µL of 0.1% TFA in water and fractionated according to Pierce High pH Reverse-Phased Peptide Fractionation kit (Thermo Scientific #84868) instructions. Fractions were dried via speed vac. Dry samples were resuspended in 25 µL 95% MS-grade water, 5% MS-grade acetonitrile and 0.1% formic acid, then 1 µg was injected on a QExactive LC-MS/MS. Raw proteomic data for the control, WT, and V122I heart tissue is presented in Table S3. Raw proteomic data for fibril extracts from WT and V122I heart tissue are presented in Table S4.

### Mass Spectrometric Identification of WT TTR Tryptic Fragments

Fibrils extracted from the WT ATTR patient heart were suspended in 8 M urea and reduced with 10 mM dithiothreitol (DTT), followed by alkylation with 25 mM iodoacetamide (IAA). The sample was subsequently diluted to a final urea concentration of 1 M using 50 mM EPPS buffer, and trypsin (Thermo Scientific Pierce, #90057) was added for overnight digestion at 37 °C. After digestion, samples were acidified with trifluoroacetic acid (TFA) and peptides were desalted using C-18 PipetteTip (Thermo Scientific Pierce, #87784) cleanup according to the manufacturer’s protocol. Eluted peptides were dried by SpeedVac and resuspended in 0.1% formic acid. For LC–MS/MS analysis, 50 ng of peptide was injected per run.

Peptide samples were analyzed on an Orbitrap Exploris 480 mass spectrometer coupled to a Vanquish Neo UHPLC system (Thermo Fisher Scientific). Peptides were first loaded onto a PepMap™ Neo Trap Cartridge (300 μm × 5 mm, C18, 5 μm, 100 Å) and subsequently separated on an EASY-Spray™ PepMap™ Neo analytical column (75 μm × 500 mm, C18, 2 μm, 100 Å). Chromatographic separation was performed at a flow rate of 350 nL/min with the column maintained at 45 °C. The initial mobile phase consisted of 98% buffer A (0.1% formic acid in water) and 2% buffer B (0.1% formic acid in 80% acetonitrile). Peptides were eluted using a linear gradient from 2% to 29% buffer B over 29 min, followed by an increase from 29% to 42% buffer B over 18 min, and then to 50% buffer B over 5 min. The column was subsequently washed with 99% buffer B for 8 min.

Mass spectrometry data were acquired in data-dependent acquisition (DDA) mode. Full MS1 scans were collected over an m/z range of 320–1600 at a resolution of 120,000 with an automatic gain control (AGC) target of 3 × 106. Precursor ions were isolated using a 3.6 m/z isolation window. MS2 scans were acquired at a resolution of 15,000, with fragmentation performed by higher-energy collisional dissociation (HCD) using a normalized collision energy of 28%. Up to 20 MS2 scans were collected per MS1 scan. Monoisotopic precursor selection and dynamic exclusion were enabled, and precursor ions with charge states from +1 to +6 were selected for fragmentation. MS1 spectra were acquired in profile mode, while MS2 spectra were acquired in centroid mode.

Raw LC-MS/MS data files were processed using FragPipe (v22.0). Peptide identification was performed with MSFragger (v4.1) using a FASTA file of full-length human transthyretin (TTR; UniProt: P02766). Default search parameters were applied except for enzyme specificity: protein digestion was set to nonspecific cleavage, with enzyme 1 defined using strict trypsin rules and enzyme 2 set to null. The precursor mass tolerance was set to 20 ppm, and the fragment mass tolerance was set to 20 ppm. Carbamidomethylation of cysteine was specified as a fixed modification, and oxidation of methionine was included as a variable modification. Label free peptide quantification was carried out using IonQuant (v1.10.27) with default parameters. Match-between-runs was disabled, and intensity normalization across runs was not applied, as only a single LC–MS/MS run was analyzed. The complete set of tryptic, semi-tryptic, and other TTR peptides detected in this experiment are presented in Table S5.

### Statistical Analysis

An average (geometric mean) normalized intensity mass-to-charge ratio was obtained across three distinct regions of each sample. Statistical analysis to determine significant differences between these ratios in diseased tissue (WT or V122I variant) compared to the control was performed on the processed mass spectrometry data using the LIMMA R package.^21^ Reported fold change ratios are ratios of the geometric means of the intensities measured from each area of the diseased hearts and the control heart. A multiple-testing adjusted p-value (p_adj_) cutoff of <0.05 was used for significance.

### Negative Staining TEM

Sample quality was assessed by negative-stain electron microscopy following established protocols.^20,22^ In brief, ATTR was applied to glow-discharged 400 square mesh copper/rhodium Maxaform grids (Electron Microscopy Sciences), incubated for 1 min to allow for fibril adsorption, and stained with 2 % uranyl formate. Grids were blotted with Whatman #1 filter paper from the side for 1 sec until dry. Micrographs were collected automatically with Leginon^22,23^ on a Thermo Fisher Talos F200C transmission electron microscope operating at 200 keV and equipped with a Gatan K2 direct electron detector at a nominal magnification of 73,000, corresponding to a pixel size of 1.981 Å (Figure S1). To study antibody binding of amyloids, a suspension of ATTR fibril-containing extract (100 μL, ∼0.5 μg/μL protein content) was incubated at a 1:1 mass ratio with anti-TTR antibodies 5D3 (Protego Biopharma) and PA-35315 (Invitrogen) and control anti-α-tubulin antibody DM1A (Cell Signaling Technology) for 1 h. Before TEM imaging, free antibody was removed by pelleting the amyloids and resuspension in antibody-free buffer.

### Preparation of Samples for Cryo-EM

UltraAuFoil 200 mesh R 2/2 grids were glow-discharged under vacuum for 30 s at 15 mA in a Pelco easiGlow 91000 glow discharge cleaning system (Ted Pella). 3.5 µl sample was applied to the front of the grid and 0.5 µl to the back, incubated for 1 min, and back-blotted with Whatman 1 filter paper for 4-6 s after the liquid spot on the filter paper stopped spreading. Grids were manually plunge-frozen in a 4 °C cold room with >95% humidity. AutoGrids were assembled and immediately screened.

### Cryo-EM Data Collection

Cryo-EM datasets were collected on a Talos Arctica TEM (Thermo Fisher) operating at 200 keV. Movies were recorded using a Falcon 4i direct electron detector (Thermo Fisher) at a nominal magnification of 150,000, corresponding to a pixel size of 0.94 Å. Movies were saved in the electron-event representation (EER) format and recorded at a total electron exposure of 50 e− per Å2. All datasets were collected automatically using EPU (v.3.9, Thermo Fisher) with a defocus range of −0.8 to −2.0 μm. EPUs fast exposure navigation was used to collect data at an 8 µm image shift. All datasets were processed using cryoSPARC (v.4.6). The processing workflow is shown in Figure S2 and described in more detail in a recent publication.^24^ Movies were collected and dose-fractioned into 40 frames, motion corrected using patch-based motion correction, followed by patch-based CTF estimation in cryo-SPARC live. Micrographs with CTF fits above 6 Å and astigmatism above 600 Å were discarded. The remaining accepted micrographs were used for further processing. Per-dataset micrograph statistics are compiled in Table S2.

### Helical Reconstruction

Filaments were picked with the filament tracer using a 60 Å diameter and a segment separation distance of 9 Å. Segments were extracted in a 300 px box, Fourier-cropped to 100 px, and subjected to 2D classification using 100 classes, and minimization of per-particle scales was enabled. Selected particles were used for initial Helical refinement without supplying helical symmetry parameters using a 90 Å diameter cylindrical starting model. A second round of helical refinement ran with 4.8 Å helical rise and - 1.3° helical twist and the previous helical reconstruction. Segments were re-extracted without binning and re-centerng turned off. Re-extracted particles were subjected to another round of helical refinement and passed into 3D classification in input mode with three or four classes and an initial lowpass of 4 Å. Reconstructions with cross-beta stack separation were combined and refined using helical refinement and a starting resolution of 5 Å. Local and global CTF refinement was performed, optimizing tilt, spherical aberration and anisotropic magnification for two iterations. A final helical reconstruction was performed with non-uniform refinement enabled, an initial low-pass resolution of 4 Å, 4.85 Å rise, -1.3° twist, a helical symmetry order of three and minimize over per-particle scale enabled. Using 171,067 segments for ATTR wt apex1 resulted in a 3.2 Å reconstruction (FSC 0.143, B-factor -46 Å^2^) with an optimized twist of -1.26° and rise of 4.85 Å. Using 342,253 segments for ATTR wt apex2 resulted in a 3.4 Å reconstruction (FSC 0.143, B-factor -68 Å^2^) with an optimized twist of -1.28° and rise of 4.85 Å. Using 335,693 segments for ATTR V122I resulted in a 3.0 Å reconstruction (FSC 0.143, B-factor -36 Å^2^) with an optimized twist of -1.24° and rise of 4.85 Å. Reported B-factors result from unsupervised auto-sharpening in RELION 5.^23,25^ A full validation of the reconstruction in shown in Figure S2.

### Reconstruction of COLVI Microfibrils

COLVI triple helical fibrils were processed in cryoSPARC (v4.7). Non-amyloid fibrillar 2D classes were selected from the processing of ATTR amyloids (wt-1) and used for template picking of previously denoised micrographs (Pretrained Model). 1.8 million picked particles were extracted in a 512px box, binned to 256px and subjected to two rounds of 2D classification (non-default parameters: 40 iterations, batch size 400, 250 Å circular mask). 147,000 particles were used to create initial reconstructions, generating three classes. All three classes were used as input heterogeneous refinement (HR; non-default parameters: hard classification enabled, 4000 batch size, spherical mask 280 Å). Finally, a set of 48,463 particles were selected from the HR and used as input for non-uniform refinement (non-default parameters: extra passes 5, initial low pass 15 Å, minimize over per-particle scale enabled), providing a final reconstruction of 9.7 Å (Gold-standard FSC resolution at 0.143). Validation and a processing workflow are shown in Figure S3 and statistics are in Table S2.

### Reconstruction of COLVI-ATTR Co-Fibrils

COLVI fibrils were processed in cryoSPARC (v4.7). Non-amyloid fibrillar 2D classes were selected from the processing of ATTR amyloids (wt-1) and used for template picking of previously denoised micrographs (Pretrained Model). 1.8 million picked particles were extracted in a 512px box, binned to 256px and subjected to two rounds of 2D classification (non-default parameters: 40 iterations, batch size 400, 250 Å circular mask). 147,000 particles were used to create initial reconstructions, generating three classes. All three classes were used as input heterogeneous refinement (HR; non-default parameters: hard classification enabled, 4000 batch size, spherical mask 280 Å). Finally, a set of 48,463 particles were selected from the HR and used as input for non-uniform refinement (non-default parameters: extra passes 5, initial low pass 15 Å, minimize over per-particle scale enabled), providing a final reconstruction of 9.7 Å (Gold-standard FSC resolution at 0.143). Validation and a processing workflow are shown in Figure S3 and statistics are in Table S2.

### Model Building, Refinement, and Validation

To aid model building, sharpened maps were modified using EMReady,^26^ which improved interpretability during model building. PDB 8G9R was used as the initial atomic model and fitted into the densities as rigid bodies within ChimeraX and manually corrected using Coot (v.0.9.8.95)^27^ and ISOLDE^28^ and iteratively refined using phenix.real_space_refine within Phenix (v.1.19) with NCS constraints enabled on 6 chains. Validation reports were automatically generated using MolProbity^29^ within Phenix. Q-scores were calculated within ChimeraX (https://github.com/tristanic/chimerax-qscore).^30^ Amyloid z-stack schematics were generated with atoms2svg.py.^31^ The full validation report is provided in Table S2.

### SDS-PAGE

TTR samples were diluted to desired concentrations in DPBS according to intensity of signal determined via pilot immunoblotting experiments. Samples were then aliquoted into 8-tube PCR strips and Laemmli SDS sample buffer (6x) was added. The tubes were briefly vortexed/centrifuged to mix. Samples were boiled at 100°C for 10 minutes and then cooled to 4°C in a PCR instrument (BioRad C1000 Touch #1851148). Tubes were again briefly centrifuged to re-incorporate any evaporated solution and then 11 µL per sample was added to a 4-20% Mini-PROTEAN® TGX™ Precast Protein Gel, 15-well, 15 µl (BioRad #4561096). For approximating the size of labeled proteins, 7µL of Precision Plus Protein™ All Blue Standards (BioRad #1610373EDU) were included in each gel. After loading reference ladder and samples, the electrophoresis cell (Mini-PROTEAN® Tetra Vertical Electrophoresis Cell for Mini Precast Gels, 4-gel #1658004) was connected to the power source (BioRad PowerPac™ Basic Power Supply #1645050), and gels were run for 5 minutes at 50 V to set protein into the gel, after which voltage was increased to 110 V and observed intermittently, removing the gel immediately after the dye front had begun to run off the bottom of the gel.

### Western Blotting

SDS-PAGE gels were transferred onto a polyvinylidene difluoride (PVDF) membrane (Sigma #IPFL00010) using a BioRad TransBlot Turbo semi-dry transfer system (BioRad #1704150) at 25 V for 7 minutes on the pre-set mixed molecular weight program. The PVDF membrane was then washed briefly with 1× tris-buffered saline solution (TBST) with 0.05% Tween20 (Sigma P7949-500 mL) and then blocked for >30 minutes in TBST+5% w/v Bovine Serum Albumin (BSA; Heat Shock Treated, Fisher BP1600-100) and 0.2% sodium azide (Fisher BP922I-500) on an orbital shaker at room temperature. After 30 minutes in blocking buffer, the buffer was removed, the membrane was washed with TBST, and polyclonal rabbit transthyretin antibody (Invitrogen Antibody, Thermo PA5-35315) was diluted 1:2000 in 5% BSA+NaN_3_ blocking buffer and added. The box was then wrapped in parafilm to preserve the primary antibody solution and placed on an orbital shaker at 4 °C overnight. The following day, the primary antibody solution was returned to a conical tube and stored at 4 °C, and the membrane was washed quickly with TBST twice followed by two additional washes with TBST on an orbital shaker at room temperature for 5 minutes each. After the fourth wash, a secondary antibody solution was added, made with TBST+3% w/v BSA and a 1:10000 dilution of HRP-conjugated rabbit antibody (Cell Signaling Technology #7074). The membrane was returned to the orbital shaker at room temperature for 1 hour, then washed again four times as described above. Depending on signal intensity, the membranes were then agitated briefly in either SuperSignal West Pico PLUS (Thermo #34580) or Femto (Thermo #34096) Chemiluminescent Substrate and then placed in a plastic sheet protector and developed on a BioRad Chemidoc using the Chemi High Resolution protocol with appropriate exposure durations. For composite images to show the Precision Plus Protein™ All Blue Standards (BioRad #1610373EDU), blots were again developed using a multichannel protocol with the second channel being Cy5 with automatic exposure set to detect intense bands.

## RESULTS

### Clinical Profile of Cardiac Tissue Samples

Two hearts with biopsy-proven ATTR CM pathology and one age-matched control heart were used for this study. Both diseased human hearts were explanted during orthotopic heart transplantation and represent end-stage ATTR CM. Upon genetic testing, the heart explanted from a 57-year-old Black female harbored only WT ATTR, hence she was diagnosed with sporadic WT-ATTR CM. The other diseased heart was explanted from a 63-year-old Black male who was diagnosed with hereditary V122I CM. The age-matched human control heart was explanted by UNOS following a non-cardiac death of a 60-year-old male of Caribbean (Puerto Rico) descent with no known cardiovascular history. Amyloid-specific staining and proteomic testing confirmed that the control heart was amyloid-free, and structural staining of the control revealed normal cardiac architecture as expected. Sample preparation from each heart occurred concurrently to mitigate experimental variability. Additional biological replicates for late-stage WT and variant ATTR CM, as well as hearts from early-stage WT and variant ATTR CM, will be sought for future studies, as at this point these results should be considered exploratory.

### ATTR CM Proteome Profile by Mass Spectrometry

Three samples from each heart (200 mg of ventricular tissue, each from a different region of the heart) were homogenized in PBS using an Ika-Werk Ultra-Turrax tissue homogenizer. Each sample was then split, with half of the samples incubated overnight in 6M guanidine thiocyanate. Mass spectrometry-based tissue analysis revealed the expected ATTR proteomic signature, i.e. elevation of serum amyloid P protein (SAP), apolipoprotein E, clusterin, vitronectin, and apolipoprotein A-IV for the diseased samples compared to the control (Figure 1A). Isobaric labeling using tandem mass tags (TMT) allows multiplexing and facilitated quantification. The intensities for each sample group were plotted in Figure 1A, and the significance of the intensity differences was determined using the LIMMA R package.^21^ The samples consisted of 3 replicates, each from a different region of the heart (see Materials and Methods). Complete proteomics results from the tissue samples are given in Table S3.

**Figure 1.**
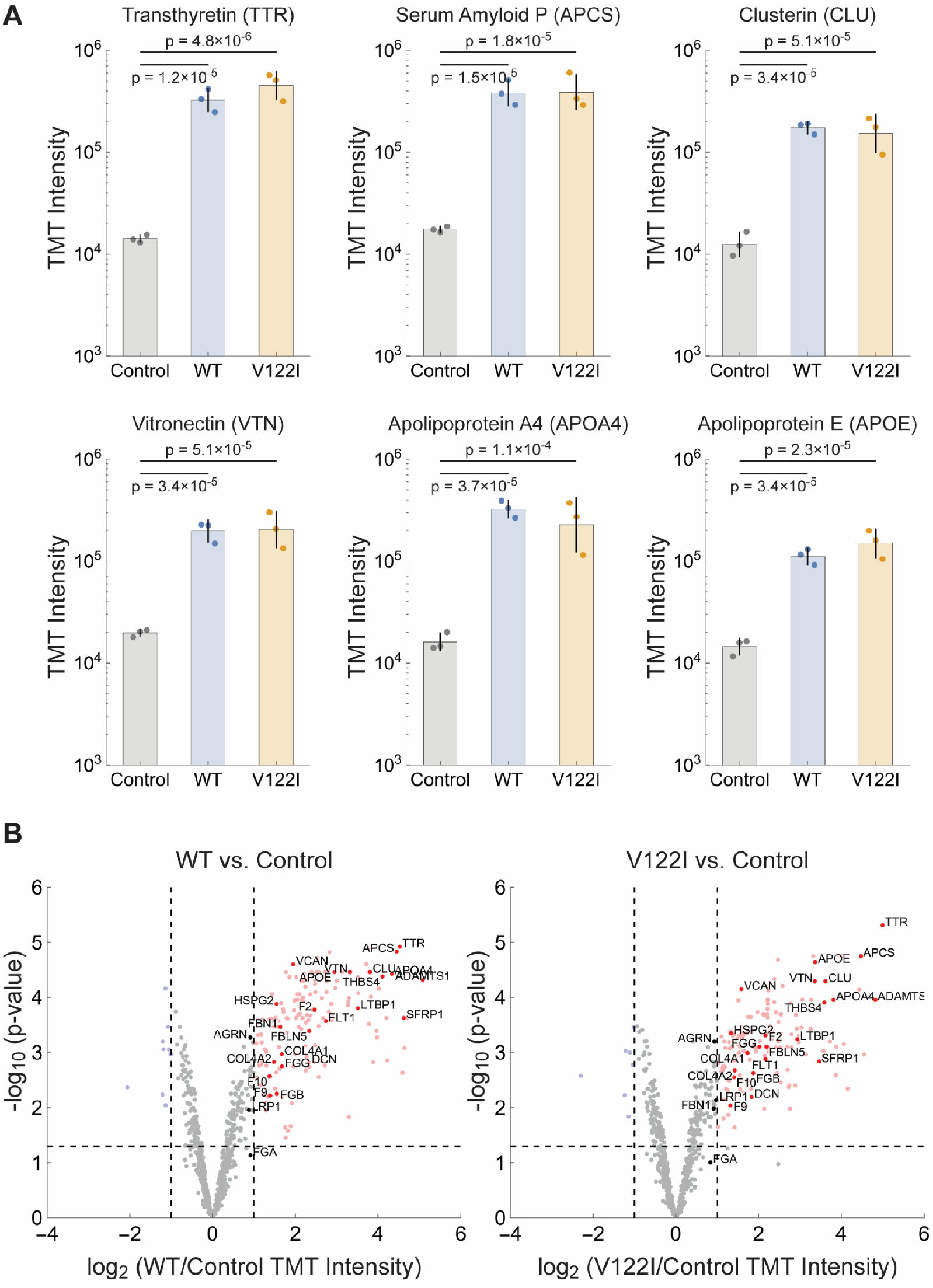
Signature Amyloid Proteins via Proteomics. A. Signature amyloid proteins identified via bottom-up proteomics of ventricular tissue homogenate from two diseased samples, WT-ATTR & V122I-ATTR, compared to an age-matched control. Bars represent the geometric means of TMT intensities of proteins in each tissue; error bars represent the geometric standard deviation. Reported p-values are adjusted p-value calculated with the LIMMA algorithm. Note the logarithmic scale on the y-axis. B. Plot of the log_2_ ratio of the TMT intensities of identified proteins in ventricular tissue homogenate from WT ATTR (left) or V122I ATTR (right) diseased tissue compared to the age-matched control (x-axis) vs. the -log_10_ adjusted p-value of the ratio calculated with the LIMMA algorithm (y-axis). The horizontal dashed line indicates p = 0.05. The vertical dashed lines indicate two-fold differences between diseased or control tissue. Data for proteins with a two-fold difference favoring the control tissue and p < 0.05 are shown in light blue. Data for proteins with a two-fold difference favoring diseased tissue are shown in light red. Data for other proteins are in gray. Proteins mentioned in the text are labeled and shown in darker colors.

Transthyretin (TTR), the protein deposited in ATTR CM, demonstrated a statistically significant fold change ratio (FCR) increase compared to the control for both WT-ATTR CM (23.0 FCR, p_adj_ = 1.2×10^−5^) and V122I ATTR CM (31.9 FCR, p_adj_ = 4.9×10^−6^). The TTR:albumin ratio in the diseased sample homogenate was 0.91 for WT-ATTR and 1.78 for V122I ATTR (Table S3), and the ratio in the extracted fibrils was 11.46 for WT ATTR and 10.01 for V122I ATTR (Table S4). We also observed statistically significant elevation of the FCR for the serum amyloid P (SAP) acute response phase protein, clusterin, vitronectin, and amyloid-associated apolipoproteins APOA4 and APOE. SAP (encoded by the APCS gene), known to bind to non-native protein conformations including amyloid fibrils, is secreted from the liver (in response to IL-6) into the blood wherein it serves as a humoral pattern recognition receptor that can directly activate complement or bind to immune cell Fc-gamma receptors to stimulate phagocytic or cytotoxic cells. SAP binds to the collagen-like region of C1q in the presence of calcium, thereby activating the classical complement pathway.^32^ SAP also plays an important role in regulating extracellular matrix composition and is co-localized with nearly all proteins having a cross-β-sheet amyloid structure.^33^ SAP demonstrated a statistically significant FCR for WT ATTR CM compared to the control of 21.9 (p_adj_ = 1.5×10^−5^) and the V122I ATTR CM FCR was 22.2 (p_adj_ = 1.8×10^−5^). Clusterin is an extracellular chaperone (ATP-independent) that binds to non-native proteins to neutralize their proteotoxicity.^34^ Clusterin also mediates non-native protein degradation by myeloid cells employing receptor-mediated endocytosis. Once internalized by macrophages or microglia (for example), complexes between CLU and misfolded proteins are trafficked to lysosomes where they are degraded by lysosomal proteases.^35^ Clusterin is elevated at an FCR of 13.9 (p_adj_ = 3.4×10^−5^) in WT ATTR CM and 12.3 (p_adj_ = 5.1×10^−5^) in V122I ATTR CM. Vitronectin, a glycoprotein secreted by the liver, seems to co-deposit with most proteins adopting a cross-β-sheet amyloid conformation^36^ and is involved in hemostasis and vascular remodeling, through interactions with integrins and extracellular matrix and blood vessel components.^37^ The amyloid fibril-associated glycoprotein vitronectin (VTN)^38^ was elevated at an FCR 10.0 (p_adj_ = 3.4×10^−5^) in WT ATTR CM and 10.3 (p_adj_ = 5.1×10^−5^) in V122I ATTR CM. Amyloid-associated apolipoproteins demonstrated a statistically significant FCR of 20.2 (p_adj_ = 3.7×10^−5^) in WT ATTR CM and 14.0 (p_adj_ = 1.1×10^−4^) for V122I ATTR CM for apolipoprotein A-IV (APOA4), and an FCR of 7.7 (p_adj_ = 3.4×10^−5^) in WT ATTR CM and 10.3 (p_adj_ = 2.3×10^−5^) in V122I ATTR CM for apolipoprotein E (APOE).

In total, mass spectrometry identified 17,393 peptides, comprising 792 unique proteins of varying statistical significance in ATTR CM in comparison to the control heart samples (Figure 1B, Table S3). The observed overall trend indicated phenotypic convergence between WT ATTR CM and V122I ATTR CM; in fact, no proteins were identified to be significantly different (p_adj_ value < 0.05) when comparing WT ATTR to V122I ATTR (Table S3).

### Coagulopathy and Complement Cascade in Cardiac ATTR Amyloidosis

Several clotting factors from the coagulation cascade were observed to be increased in the ATTR CM tissue homogenate compared to the control (Figure 2). Most of the components of fibrinogen (clotting factor I) were increased with the β-chain (FGB) exhibiting an FCR of 2.9 (p_adj_ = 5.6×10^−3^) in WT ATTR CM and 3.7 (p_adj_ = 2.4×10^−3^) in V122I ATTR CM, and a γ-chain (FGG) FCR of 3.2 (p_adj_ = 1.8×10^−3^) in WT ATTR CM and 4.1 (p_adj_ = 7.8×10^−4^) in V122I ATTR CM (the α-chain (FGA) was present at higher levels in the ATTR hearts but did not quite reach significance, with an FCR of 1.9 (p = 0.073) in WT ATTR CM and 1.8 (p = 0.098) for V122I ATTR CM). Prothrombin (F2) was increased, yielding an FCR of 5.5 (p_adj_ = 1.7×10^−4^) in WT ATTR CM and 4.5 (p_adj_ = 4.9×10^−4^) in V122I ATTR CM. Increases in clotting factors IX (F9) and X (F10) were also observed, with an FCR of 2.6 (p_adj_ = 6.0×10^−3^) and 2.6 (p_adj_ = 2.7×10^−3^) in WT ATTR CM and 2.5 (p_adj_ = 9.1×10^−3^) and 2.7 (p_adj_ = 2.8×10^−3^) in V122I ATTR CM, respectively. In addition, in tissue homogenate, we note that proteomic analysis of the extracted fibrils from the WT and V122I ATTR hearts similarly reveals components of the extrinsic coagulation cascade as well as the classical complement cascade (Figure 2). Also present are von Willebrand factor and related proteins, platelet-related proteins, and the kinin-kallikrein system proteins (Table S3).

**Figure 2.**
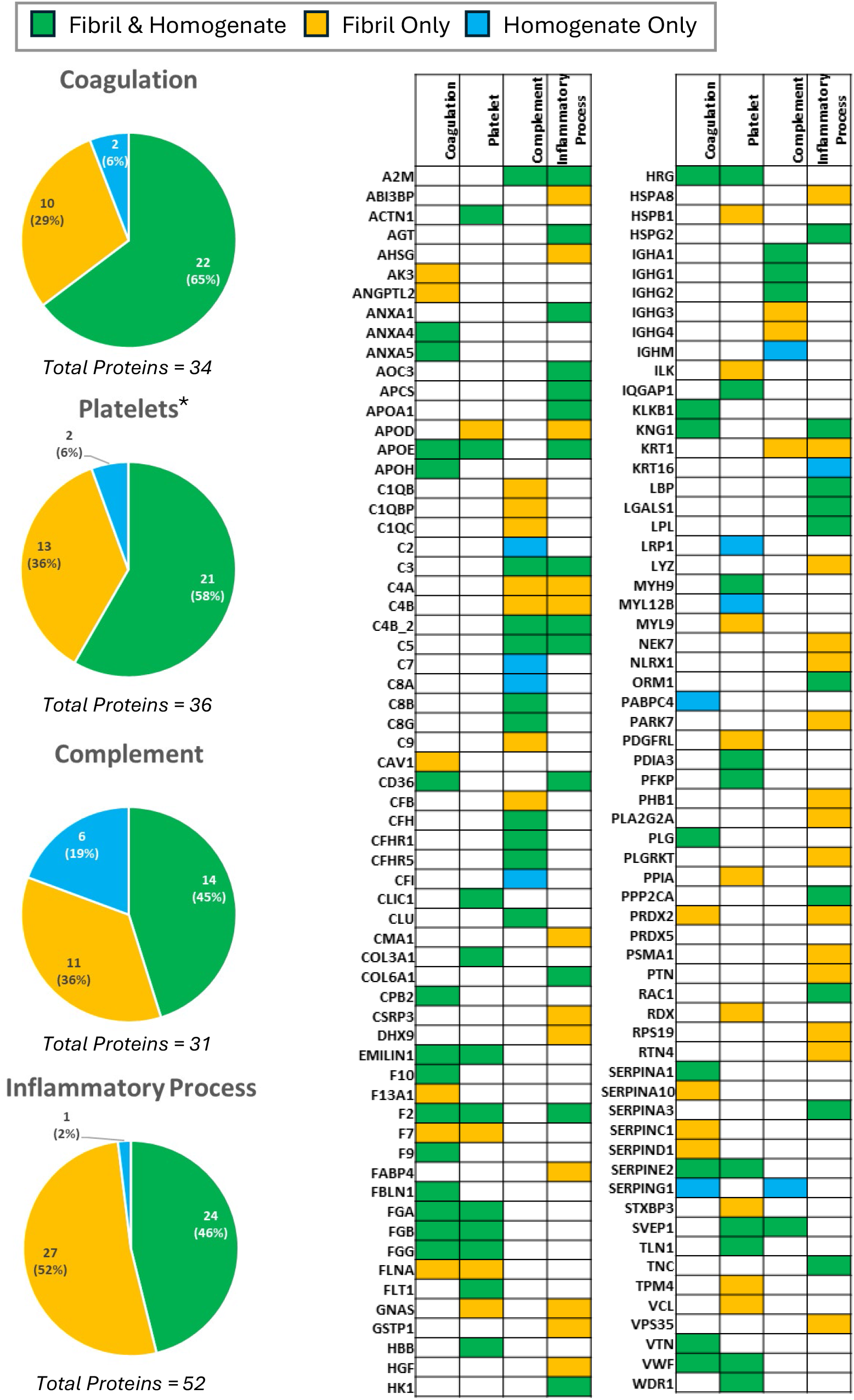
Thromboinflammatory Proteomic Profile. Proteins detected at higher levels in the ventricular tissue homogenate of patients with ATTR CM relative to control, extracted amyloid fibrils of the same patients, or both, as identified in the Gene Ontology Biological Processes database. *Platelets proteins include those pertaining to platelet aggregation, formation, activation, maturation, regulation, platelet morphogenesis, and platelet-derived growth factor.

### 3D Tissue Analysis Reveals Microvascular Occlusion via Capillary Thrombosis in Cardiac ATTR Amyloidosis

Optically cleared samples were labeled for vascular endothelial-cadherin (CD144; endothelial cell gap junctions) and thrombin via indirect immunofluorescence. These samples were also treated with AmyTracker, an oligothiophene dye that becomes fluorescent upon binding to amyloid fibrils (Ebba Biotech). 3D visualization (Figure 3) reveals the presence of thrombin in vessels measuring < 10 µm in diameter, corresponding with vascular thrombosis at the level of the capillary. Loss of vascular integrity is observed downstream of the thrombus (Figure 3) a location corresponding to the margins of amyloid staining. Loss of vascular architecture downstream of the clot indicates capillary rarefaction, with angiogenic revascularization required to restore full functionality to the underlying cardiomyocytes.

**Figure 3.**
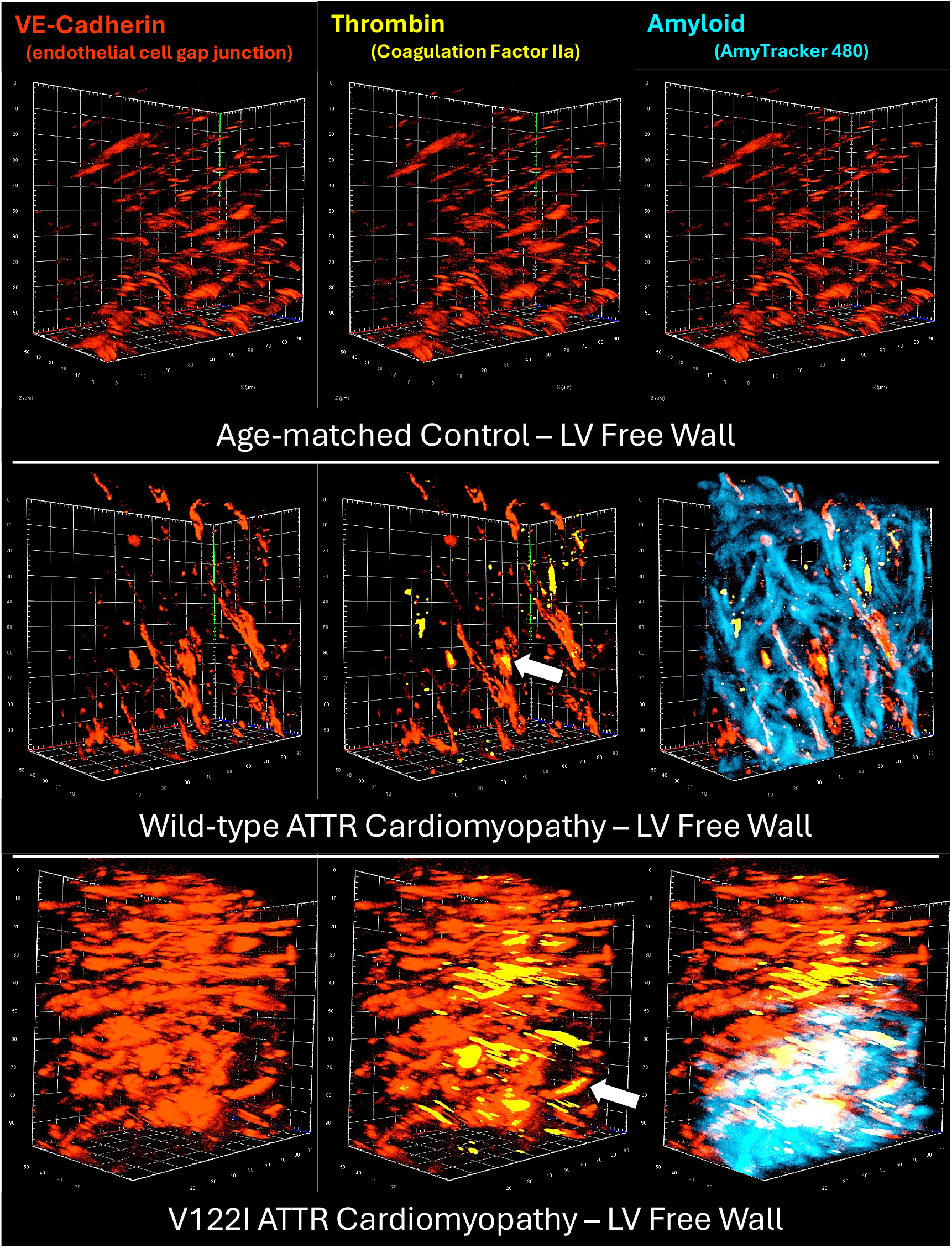
3D Visualization of Microvascular Thrombosis. Optically-cleared left ventricular free wall tissue reveals clotting (thrombin, yellow) within blood vessels of <10 μm in diameter (white arrow), signifying microvascular thrombosis of the capillaries. Occluded capillaries were frequently observed proximal to amyloid deposits, although a formal quantitative analysis was not conducted due to the small number of patients involved in the study.

### Elevated Proteins/Proteoglycans Associated with Angiogenesis

Numerous proteins and proteoglycans associated with angiogenesis are elevated in the diseased samples (Figure 4; Table S3). Collagen IV is the most abundant protein in vascular basement membranes, and collagen IV-α1 chain (COL4A1) and collagen IV-α2 chain (COL4A2) are increased, displaying an FCR of 3.20 (p_adj_ = 1.1×10^−3^) and 2.8 (p_adj_ = 1.4×10^−3^) for WT ATTR CM and 3.3 (p_adj_ = 1.0×10^−3^) and 2.7 (p_adj_ = 2.1×10^−3^) in V122I ATTR CM, respectively. Basement membrane component perlecan (HSPG2), a sulfonated proteoglycan that promotes TTR aggregation,^39^ is increased by an FCR of 2.9 (p_adj_ = 1.3×10^−4^) in WT ATTR CM disease and 2.5 (p_adj_ = 4.4×10^−4^) in V122I ATTR CM. Other proteoglycans associated with the vascular basement membrane that participate in angiogenesis include decorin (DCN), versican (VCAN), and agrin (AGRN), which are elevated, displaying FCRs of 5.0 (p_adj_ = 1.5×10^−3^), 3.9 (p_adj_ = 2.5×10^−5^), and 1.9 (p_adj_ = 5.3×10^−4^) in WT ATTR CM and 3.6 (p_adj_ = 6.4×10^−3^), 3.0 (p_adj_ = 7.0×10^−5^), and 1.9 (p_adj_ = 6.3×10^−4^) in V122I ATTR CM, respectively (Table S3). The receptor for vascular endothelial growth factor, FLT1 is increased by an FCR of 6.7 (p_adj_ = 2.7×10^−4^) for WT ATTR CM and an FCR of 4.5 (p_adj_ = 1.3×10^−3^) for V122I ATTR CM. The receptor LRP1 also exhibited a statistically significant increased FCR of 1.8 (p = 0.011) for WT-ATTR CM and a FCR of 2.0 (p_adj_ = 7.3×10^−3^) V122I ATTR CM. LRP1 plays a critical role in organized angiogenesis, binds ApoE and amyloid-disease-associated proteins like α-synuclein and tau,^12,13^ and is upregulated by the natively folded TTR tetramer.^40^ Additional proteins associated with angiogenesis that are increased in ATTR CM include latent TGFβ-binding protein (LTBP1), thrombospondin-4 (THBS4), secreted frizzle-related protein 1 (SFRP1), ADAMTS1, fibrillin-1 (FBN1), fibulin-5 (FBLN5), other thrombospondins and ADAMTS proteins, and other angiogenesis-related proteins (Figure 4, Table S3).

**Figure 4.**
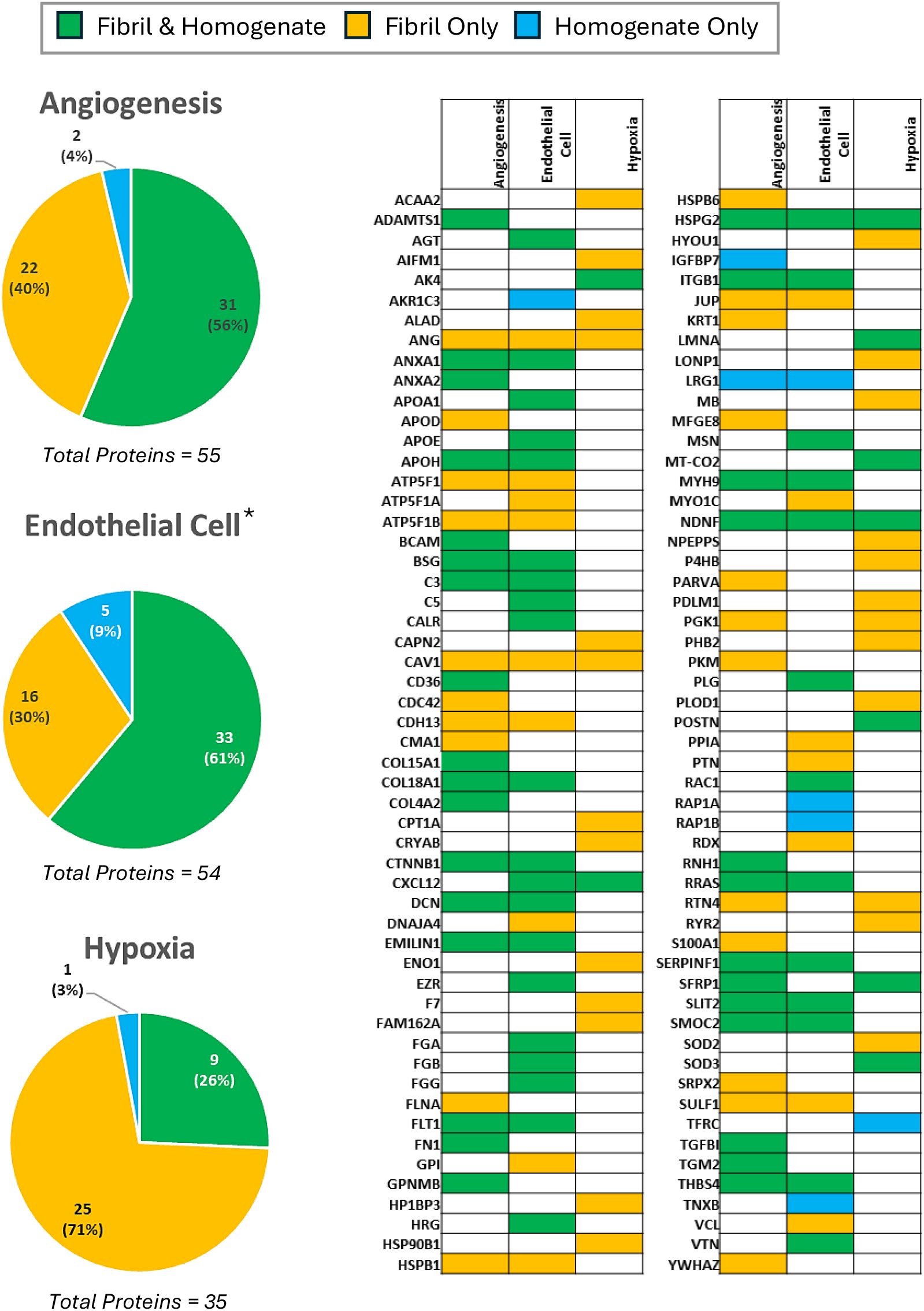
Angiogenic Proteomic Profile. Proteins identified in the ventricular tissue homogenate of patients with ATTR CM, extracted amyloid fibrils of the same patients, or both, as identified in the Gene Ontology Biological Processes database. *Endothelial Cell proteins include those pertaining to endothelial cell proliferation, migration, differentiation, regulation, morphogenesis, establishment of endothelial barrier, endothelial apoptotic process, and vascular endothelial growth factor.

### 3D Tissue Analysis Reveals Disruption of Vascular Architecture

Optically cleared samples were labeled by AmyTracker, to reveal amyloid deposits; using indirect immunofluorescence to image collagen IV, the most abundant protein in the vascular basement membrane; and by a TTR antibody (5D3) raised to a monomeric TTR construct harboring two mutations (F87M/L110M) that prevent it from forming the native TTR tetramer and consequently make it prone to slow aggregation under physiological conditions (Protego Biopharma).^41^ This antibody recognizes partially heat denatured TTR and patient-derived ATTR fibrils (Figure S4A-D) but does not bind to native tetrameric TTR, despite its sequence epitope (TSESGELHGLTTE sequence epitope, residues 49-61) being exposed (Figure S4E).^41^ These observations suggest that the binding of 5D3 depends on the conformation of its epitope, and that it can recognize this sequence broadly in non-native conformations.

Three-dimensional visualization (Figure 5; LV free wall) reveals a disruption of vascular architecture in the ATTR CM samples (bottom two rows) compared to the lattice-like architecture observed in the age-matched non-CM control LV slice images in the top row. Patches of hypervascularization (black arrows) and hypovascularization (white arrows) are observed in the cardiomyopathy sections. Amyloid deposition correlates with hypovascularized regions, suggesting an association with amyloid deposition and loss of vascularization. Interestingly, non-native TTR visualized with the 5D3 antibody overlaps only to a limited extent with fibrillar TTR deposits visualized with AmyTracker. Instead, staining with 5D3 is observed filling in areas where there is a loss of collagen IV (yellow arrows), sometimes in the absence of AmyTracker staining but more commonly along the borders of areas of dense AmyTracker staining. This observation is interpreted further in the Discussion section.

**Figure 5.**
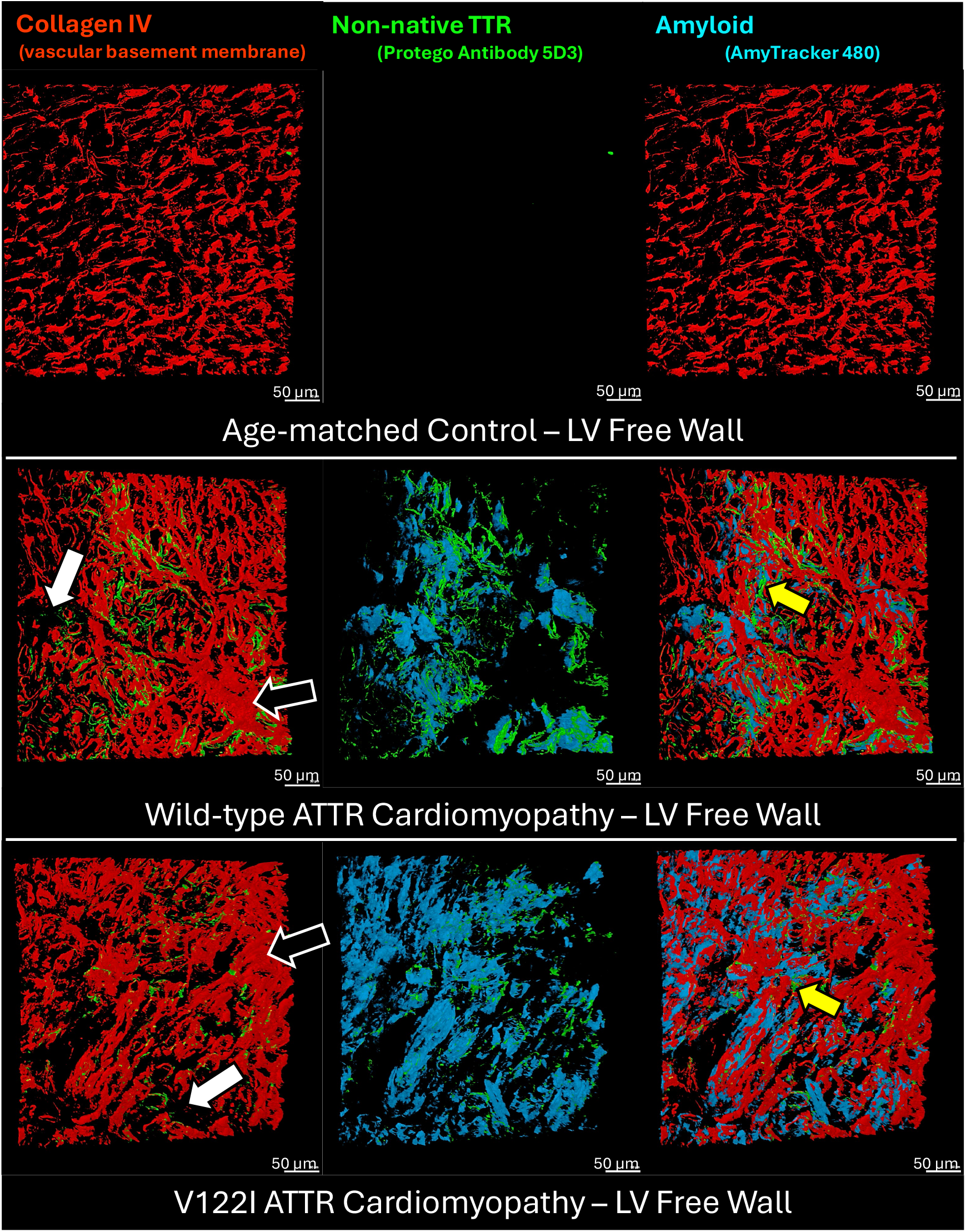
3D Visualization of Disorganized Angiogenesis. Optically-cleared left ventricular free wall tissue reveals disorganized angiogenesis. The organized lattice vascular structure observed in the age-matched control via Collagen IV staining (red) is lost in favor of hypervascular regions (black arrows) and hypovascular regions (white arrows). Non-native TTR (green) appears to replace collagen IV (yellow arrows) on the border of amyloid-rich hypovascular regions. Amyloid is stained with AmyTracker 480 (blue). Protego’s 5D3 antibody targets a TTR sequence (TSESGELHGLTTE) that is covered in its native tetramer conformation, thereby only binding to non-native forms of TTR. Non-native TTR overlaps with AmyTracker 480 on the margins of amyloid deposits, suggesting a dense fibrillar core (blue) surrounded by nonfibrillar non-native TTR aggregates (green).

### Characterization of Ex Vivo ATTR Amyloid Fibrils

To demonstrate that there were true amyloid fibrils in the WT ATTR-CM and V122I ATTR-CM hearts, we examined the extracted fibrillar material by negative stain TEM, revealing amyloid fibrils ranging widely in length but of uniform thickness (Figure S1). A left-handed helical organization was assumed but not verified experimentally. The extracted fibrils were sufficiently regular to determine their atomic structures by cryo-electron microscopy (cryo-EM). In cryo-EM, the fibrils tended to cluster, resulting in a high fraction of unsuitable micrographs (Figure S1 and S2A-C). However, enough curated micrographs were obtained that they could be used to perform automated filament tracing and helical reconstruction in cryoSPARC (Figure S2D).^24^ Cross-β sheet separation during 2D classification and 3D classification was used to select suitable homogenous stacks of helical segments to achieve reconstructions of ∼3.2 Å and ∼3.4 Å resolutions for WT ATTR fibrils and ∼3.0 Å resolution for V122I ATTR (Figure 6A-C, Figure S2). All three reconstructions showed only slight differences in helical symmetry, with a consistent rise of 4.85 Å and a twist of -1.26 ± 0.02°. Consequently, the calculated crossover distance (CO) is within a range of 682-704 Å (Figure 6D, Table S2).

**Figure 6.**
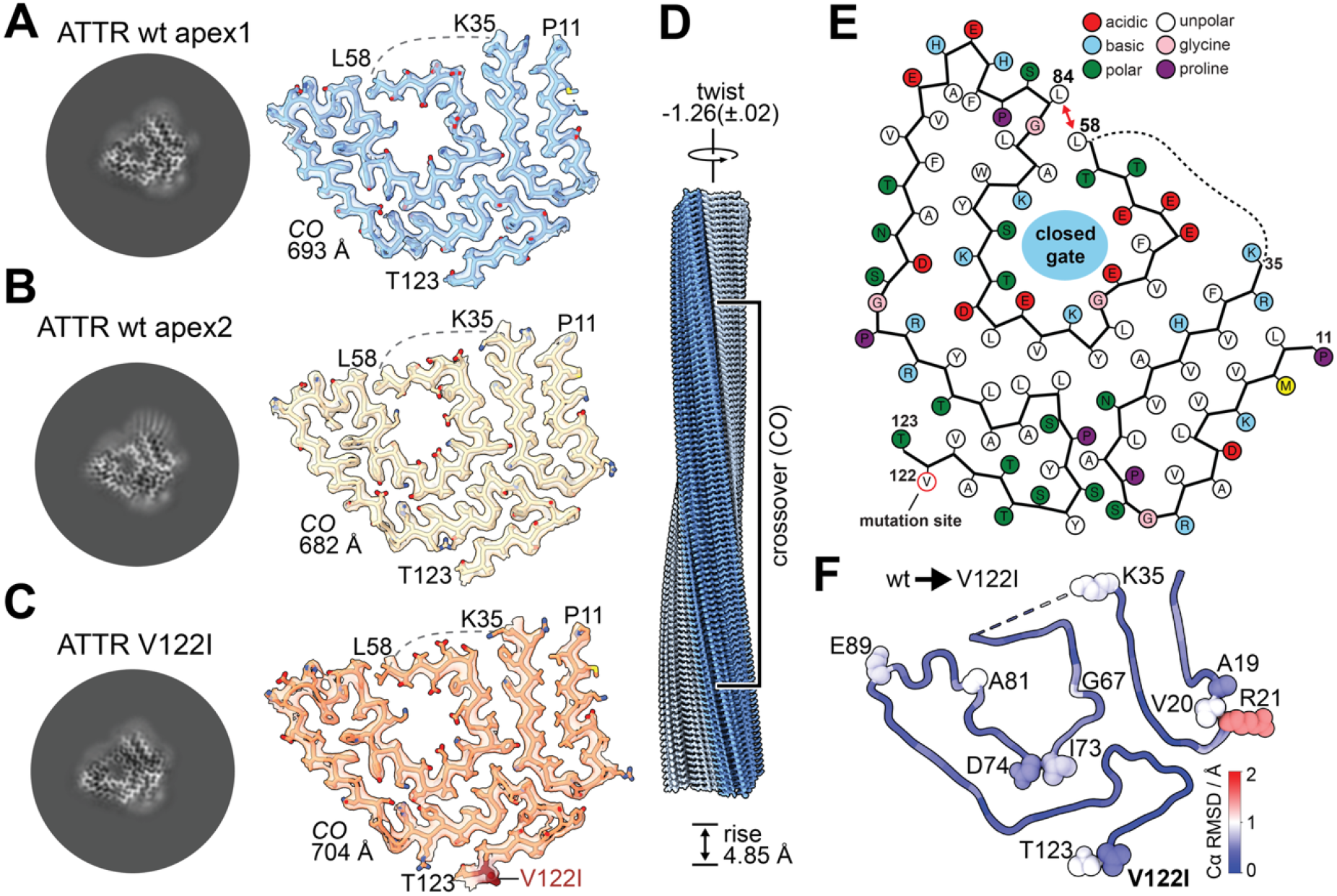
Helical Reconstruction and Characterization of Cardiac ATTR filaments. (A-C) Slices of the reconstructed ATTR filaments viewed along the Z-axis (left) shown next to an isosurface of the EM density with the atomic model within. Wild-type apex1 in blue, WT apex2 in tan, and familiar CR-variant V122I in orange. Peptide termini are annotated. (D) Consensus atomic model of the ATTR amyloid, with the small P11-K35 peptide colored blue and the large peptide L58-T123 colored grey. The helical symmetry parameters are annotated: 4.85 Å rise, -1.26(±.02)° twist and crossover (CO) with per-sample values next to the map-model fits. (E) Residue-level schematic of the consensus ATTR spearhead conformation. The closed gate between L58/L84 is marked with a red arrow. Acidic residues are colored red, basic residues blue, polar residues green, hydrophobic residues white, glycine pink, and proline violet. The dashed line indicates an unresolved residue range. V122 highlighted in red. (F) Structural conservation between wt and V122I ATTR represented as Cα RMSD (root-mean square deviation). Residues with higher RMSD values are shown as annotated spheres.

All three reconstructions match the “spearhead” conformation characterized by a closed “gate” between L58 and L85, which has been documented for the majority of resolved ATTR amyloids (Figure 6E, Figure S5A),^19,42-48^ except for polymorphs extracted from the vitreous body of the eye^43^ and cardiac I84S fibrils (Figure S5A).^45^ As shown for previous reconstructions of ATTR (Figure S5A), the cross-β structure is well-resolved for a shorter N-terminal fragment, showing density for P11-K35, and a longer C-terminal peptide with continuous density for L58-T123 (Figure 6E). The intervening segment, A36 to G57, is poorly resolved.

Comparing the three resolved structures using Cα root-mean square deviation (RMSD) shows that the overall spearhead fold is highly stable (global RMSD 0.4 Å) with minor localized differences in loop-regions of the peptides. These regions include the peptide termini and the residues A19-R21, but the point mutation V122I does not induce observable changes to the local fold (Fig. 6F). Among all resolved ATTR amyloids, the overall Cα RMSD is less than 1 Å, with the greatest difference localized to the gate region. This high stability further supports the favored spearhead conformation over other polymorphs (Figure S5B).

The spearhead fold of ATTR is commonly modelled as two discontinuous peptide chains (connected by a dashed line in Figure 6). Since helical reconstruction relies on averaging of overlapping segments and correctly applied helical symmetry, symmetrization artifacts and averaging can bias the reconstruction towards the most abundant reconstruction.^49^ Low-abundant conformational states and flexible regions are unlikely to be resolved, and their interpretation requires additional experimental effort to understand localized mutations and their disease relevance. While we could not resolve the region of A36-G57, applying a two-sigma Gaussian to the cryo-EM density shows a potentially connected but flexible region (Figure S6A). Revisiting previously resolved ATTR amyloids from the heart^42,45^ and the vitreous body of the eye^43^ show similar signs of flexible density in the region of A36-G57 (Figure S6B).

ATTR fibrils are often found to be a mixture of full-length and proteolytically cleaved peptide chains, with the cleavage site being in this flexible region. The resulting mixed full length and split-chain fibrils have been termed type A fibrils (the less common type B fibrils consist entirely of full length TTR).^19,44-46,50-53^ Prior work strongly suggests that amyloid fibril formation precedes proteolysis (although this order of events has not been conclusively demonstrated).^19,44,46,48^ After the initial proteolysis event, the resulting fragments can be trimmed further by aminopeptidases. As a result, the C-terminal fragment in type A fibrils is “ragged”, with N-termini at positions 44 to 58.^19,44,45,50^ Western blots of the isolated fibrils from the two ATTR hearts in this study employing an antibody (PA5-35315) to part of the TTR sequence (residues 71-98) that should be retained in the expected C-terminal TTR fragment revealed that in both the WT and V122I hearts, the fibrils consisted primarily of full length TTR with proteolyzed TTR as a minor component, identifying these fibrils as type A (Figure S7A). Supporting this observation, bottom-up proteomics on tryptic digests of the ATTR fibrils from the WT patient reveals fragments covering the whole sequence of TTR (Figure S7B, C).

### Cardiac ATTR amyloid fibrils are decorated with Collagen VI

Inspection of our amyloid samples reveals the presence of an unknown amyloid fibril-associated component in almost all micrographs across the three resolved ex vivo amyloid extracts. These ATTR fibrils with their additional component adopt a repeating arrangement, owing to an apparent ATTR-interacting macromolecule associated with the ATTR amyloid fibrils, that itself exhibits a fibril morphology (Figure 7A, B). In earlier work with cardiac light-chain amyloids, similar assemblies were identified as AL amyloid fibrils interacting with collagen VI (COLVI), affording semi-structured super-helical assemblies.^54^ COLVI links the cell surface to the extracellular matrix, conferring mechanical strength to mammalian tissues. It is ubiquitously expressed, forming triple-helical fibrils with the most abundant repeating subunit-configuration being α1-α2-α3 heterotrimers,^55^ which was characterized using cryo-EM recently.^56^ We used single-particle analysis and reconstructed the single-bead portion of the non-amyloid COLVI microfibrils from our WT apex1 patient heart dataset to about 10 Å without enforcing point-group symmetry (C1) (Figure S3). All COLVI subunits include proline-rich collagen triple-helices and repeating von-Willebrand-factor-type-A (vWA) globular domains, which we placed into our microfibril reconstruction for visualization (Figure 7C).

**Figure 7.**
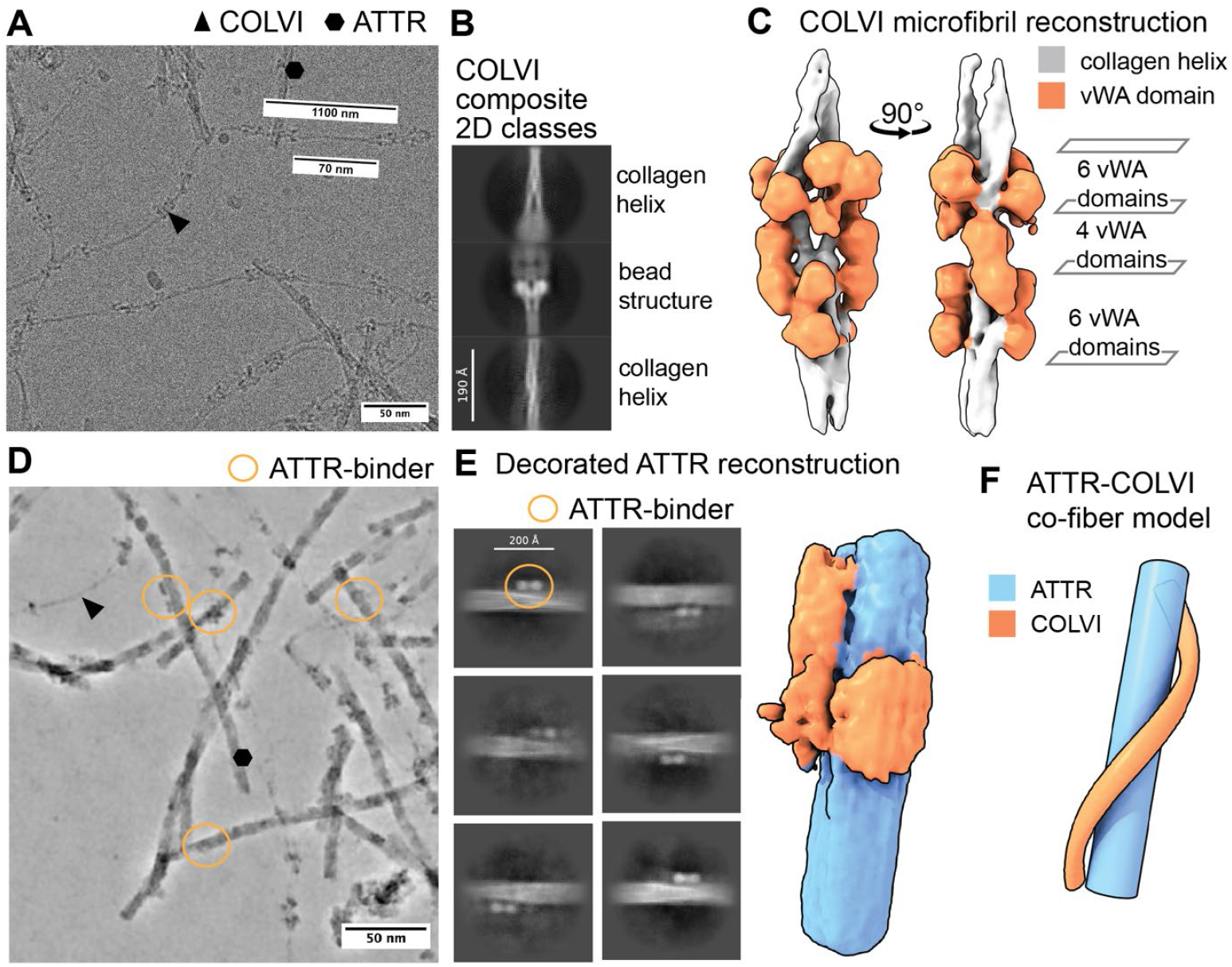
ATTR amyloid fibrils are enriched with collagen VI (COLVI). **(A)**Micrograph of cardiac ATTR amyloids (marked with black hexagon) and COLVI triple helix-based micrfibril (black triangle). **(B)** Composite 2D class average of COLVI single-bead microfibril, showing a repeating pattern of collagen helices and bead structure. **(C)** COLVI microfibril reconstruction at 10 Å, color-coded by collagen helices (light grey) and von-Willebrand-factor-type-A (vWA) domains (orange). Segments and matching numbers of vWA domains along the z-axis of the COLVI microfibril. **(D)** Denoised micrograph with regions of putative ATTR-binders (orange). ATTR amyloids (marked with black hexagon) and COLVI (black triangle). **(E)** 2D class averages of decorated ATTR amyloids and an about 10 Å reconstruction. **(F)** Model of ATTR amyloid fibril interacting with collagen VI also adopting a microfibril morphology (orange) consisting of triple helices with periodic vWA globular domains (orange)

While COLVI was not visible in the helical reconstructions of ATTR amyloids due to the applied helical symmetry, we observed non-amyloid densities decorating the ATTR fibrils on a micrograph- and projection-level (Figure 7D,E). Selecting 2D classes positive for amyloid binders resulted in a low-resolution reconstruction of decorated ATTR fibrils (Figure 7E). Limited by the given resolution, we cannot unambiguously assign this binding partner of ATTR to be COLVI. However, the fits of the COLVI domains into our observed density supports the notion that this ATTR-bound species is COLVI, as does the bead-to-bead distance of 1100 Å of the microfibrils (Figure 7A),^55,56^ and the observation that COLVI is the most abundant type of collagen found in our mass spectrometric analysis of fibril extracts (Table S4). Thus, we tentatively assigned the binding partner of the ATTR fibrils to be COLVI.

Micrograph-level measurements of the triple-helical spacer between the beads of the COLVI microfibrils (globular von-Willebrand-factor-type-A (vWA) domains) was determined to about 700 Å (Figure 7A), matching the amyloid cross-over distances for ATTR amyloids (682-704 Å), and half of the resolved AL59 light-chain amyloid crossover (1200 Å)^54^ potentially supporting co-filament formation by a process of guided chirality transfer, in which the natural amyloid twist transfers onto non-twisting COLVI triple helical fibrils. A co-fibril model of ATTR and COLVI is proposed in Figure 7F. Strong, intimate associations between extracellular matrix components and ATTR fibrils like those observed in Figure 7 demonstrate how extracellular matrix components can create tissue microenvironments that could promote the deposition of cross-β-sheet amyloid fibrils with complementary structures and/or prevent their clearance.

## Discussion

Based on the observations covered in the Results section, we hypothesize that WT and hereditary ATTR CM is a microangiopathy of dysregulated angiogenesis in the setting of thrombo-inflammatory capillary occlusion (Figure 8). This hypothesis encompasses the following sequence of events. Endothelial cell activation in the capillary beds, possibly initiated by oxidized lipoproteins, thrombin, or misfolded TTR, initiates release of tissue factor (TF) and activation of the extrinsic coagulation cascade affording a fibrin clot in a capillary (Figure 8, panel 1). Simultaneously, kallikrein converts kininogen to bradykinin, initiating vasodilation and exposure of the vascular basement membrane (VBM). Exposed collagen IV, along with serum amyloid P, triggers the synthesis of C1 complex initiating the classical complement pathway (Figure 8, panel 2). Capillary occlusion (Figure 3) triggers downstream hypoxia (Figure 8, panel 3) and requires revascularization to restore oxygenation (Figure 8, panel 4). Recent studies postulate a potential role for tetrameric TTR in angiogenesis (Table S6).^10,15^ The exposed VBM is comprised of “signature amyloid-interacting proteins”^38^ and other components, such as proteoglycans, which are known to bind misfolded TTR, creating nucleation points for amyloid fibril formation (Figure 8, panel 3; see next paragraph), possibly by conformational conversion.^39,57,58^ Without attenuation of the hypoxic signal, TTR will continue to arrive at the site, resulting in TTR accumulation and congestion of the basement membrane (Figure 8, panel 4). Basement membrane congestion limits the space for sprouting angiogenesis to occur, resulting in loss of vascular architecture visualized by areas of hypervascularization adjacent to amyloid deposits and hypovascularization in areas with amyloid deposition (Figure 5). While TF is not present in our proteomics data, a recent study in which cultured endothelial cells were treated with plasma +/-tafamidis demonstrated reduced TF expression and activity in the treated group, elucidating a role for stabilized transthyretin in clotting hemostasis.^59^ If circulating non-native TTR is responsible for the endothelial cell activation that initiates coagulation, this could explain why tafamidis is so effective in slowing CM progression in the absence of amyloid clearance.

**Figure 8.**
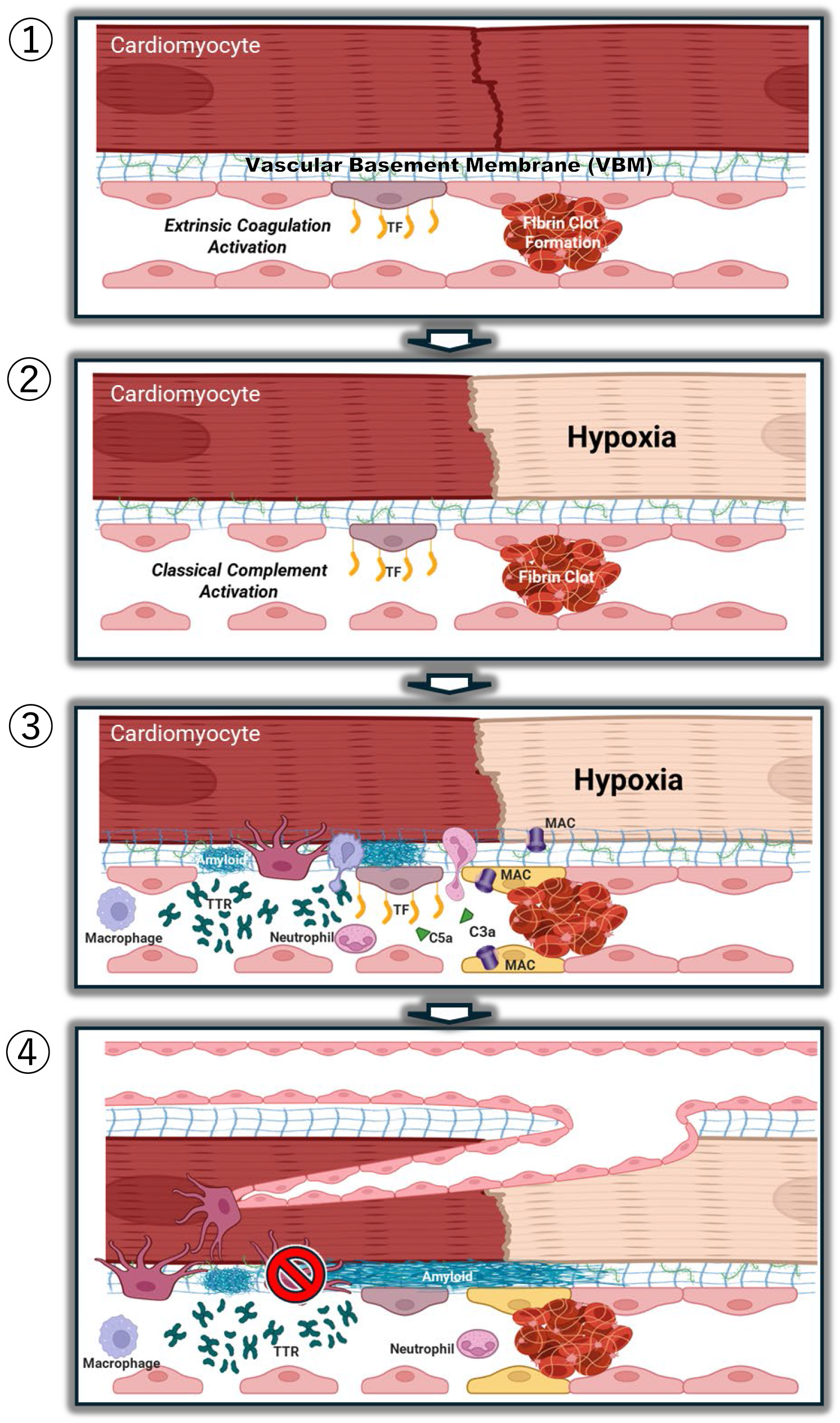
Microangiopathy and Disorganized Angiogenesis. (1) Endothelial cell activation, perhaps through contact with misfolded TTR, in the coronary capillaries leads to tissue factor (TF) expression on the luminal surface of the endothelial cell, activating the extrinsic coagulation cascade and resulting in capillary occlusion by fibrin clot formation. (2) Localized hypoxia results in an inflammatory reaction, leading to vasodilation and separation of capillary endothelial cells. Vasodilation exposes collagen IV in the basement membrane, which along with serum amyloid P activates the classical complement cascade. (3) Complement anaphylatoxins C3a and C5a recruit macrophages and neutrophils, generating reactive oxygen species and releasing proteases. Complement’s membrane attack complex (MAC) embeds in the plasma membrane of stressed cells, promoting apoptosis. TTR arrives at the site to help facilitate angiogenesis and revascularization, but misfolded TTR nucleates on exposed basement membrane proteoglycans to form amyloid fibrils. (4) Thickening basement membrane, driven by amyloid deposition, prevents efficient and organized revascularization. The new blood vessels must circumvent amyloid deposits, resulting in disorganized angiogenesis and torturous revascularization.

The ATTR-CM hearts used in this study are from patients with late-stage disease, and as such are of limited value in establishing the order of events leading to the images herein. That said, we noted above that staining mediated by the anti-non-native TTR antibody (5D3) occurs in the absence of or, more commonly, in the periphery of AmyTracker-stained amyloid deposits. Given that we showed above that the anti-non-native TTR antibody binds to isolated ATTR fibrils (Figure S4), we speculate the reason that it does not bind to the dense core of ATTR amyloid deposits likely has to do with steric occlusion of its epitope because of interactions between ATTR fibrils (consistent with their propensity to cluster in Figure S1) or between ATTR fibrils and amyloid binding proteins / glycosaminoglycans, whether circulating (like SAP) or in the extracellular matrix (like COLVI). The strength of such interactions is demonstrated by the intimate and structurally specific association between COLVI and ATTR fibrils shown in Figure 7. The anti-non-native TTR antibody is therefore likely associated with immature, non-fibrillar ATTR deposits (which are not expected to stain with AmyTracker^60^), as well as the fibrillar and non-fibrillar material around the periphery of dense amyloid deposits. Based on these observations, we hypothesize that our results are consistent with a process in which non-fibrillar TTR aggregates (oligomers / protofibrils), most likely consisting of full-length TTR, deposit first in the vascular basement membrane. Such oligomers are known to circulate in the blood in both ATTR polyneuropathy and cardiomyopathy.^41,61-64^ Two processes would then occur, likely in parallel. First, the deposited aggregate would present a favorable “landing zone” for circulating non-native oligomeric TTR because of the homotypic interactions between the two leading to growth of the deposit. Second, the deposited structurally heterogeneous aggregates would undergo conformational conversion to form denser, more structured, and more stable cross-β-sheet amyloid fibrils. Interactions with species like COLVI (Figure 7) and sulfated glycosaminoglycans^39,57,58^ in the extracellular matrix could potentially promote such a conformational conversion.

Conformational conversion processes like the one described above have been widely observed for many amyloidogenic proteins in vitro.^65-69^ Moreover, this sequence of events is consistent with observations in transgenic mouse models of ATTR, in which non-amyloid TTR deposits were shown to occur in a given organ at younger ages than fibrillar TTR deposits.^57,70^ These processes would produce TTR deposits with a dense, fibrillar core consisting of older deposited proteins associated with extracellular matrix components and other fibril-binding proteins, surrounded by a corona of non-fibrillar TTR aggregates, with the vascular pathology being initiated as soon as the initial deposits form, but persisting and worsening as the deposit grows and stabilizes as a result of the conformational conversion to a fibrillar form. In this context, it is worth noting that interactions between amyloid fibrils and COLVI^71^ (like other signature amyloid binding proteins^72-74^) have been suggested to prevent the recognition and clearance of fibrils by the immune system and could therefore impact the trajectory of ATTR cardiomyopathy at the clearance as well as at the deposition stage.

TTR’s role as an acute phase reactant in angiogenesis is still being elucidated, so we investigated other potential explanations for TTR’s abundance in the diseased tissue. Proteomics from both the tissue homogenate and extracted fibrils failed to reveal the presence of RBP4, making retinol transport an unlikely mechanism. TTR is a secondary carrier of thyroxine, and the lack of its primary carrier (TBG) in the proteomics suggests that TTR’s presence is unrelated to thyroxine delivery. Passive vascular leakage was also ruled out due to the TTR:albumin ratio.

As shown in Figure 5, the microvasculature of cardiac ATTR amyloidosis in the LV free wall is marked by loss of vascular architecture compared to that of the age-matched control, exhibiting areas of hypervascularization and hypovascularization. This is indicative of abnormal angiogenesis. Abnormal angiogenesis is often associated with a tumor microenvironment,^75^ driven by the increased metabolic needs of the growing tumor. The abnormal angiogenesis observed in our samples is unlikely to be due to increased metabolic need, as there is no tumor present, and a much more uniform distribution of microvasculature would be expected if the angiogenesis was driven by increased myocardial demand. Without an increased metabolic need, an alternative explanation for the angiogenic stimulus is local ischemic signaling due to hypoxia and loss of vascularization. Indeed, Figure 3 reveals thrombus at the level of the capillary in the presence of a coagulopathic environment (Figure 2), which would create localized hypoxia and a need for re-anastomosis – triggering angiogenesis.

While this is the first study identifying capillary thrombosis in ATTR CM, Yeh et al.^76^ recently identified microangiopathy with thromboinflammation in late-stage TTR polyneuropathy in the sural nerves of A97S ATTR variant patients. This aligns with the pathophysiology we observe in ATTR CM samples, suggesting a common pathophysiological mechanism driving both tissue tropisms. Studies performed by the Bellotti group^77^ also suggest a hemostatic pathophysiology to ATTR CM, demonstrating cleavage of TTR tetramer by both trypsin and the physiologically-relevant plasma protease plasmin – a hypothesis that was not investigated in this study.

Multiple clinical studies corroborate our findings of microvascular dysfunction in ATTR CM. In a study of cardiac amyloidosis patients (both ATTR and AL) undergoing quantitative stress perfusion cardiac MRI with myocardial blood flow mapping,^78^ Chacko et al. demonstrated that cardiac amyloidosis is associated with severe inducible myocardial ischemia characterized by diffuse hypoxia and disruption of microvascular architecture in the presence of vascular amyloid deposits, luminal stenosis, and extensive capillary loss. Netti et al. demonstrated quantifiable microvascular obstruction by cardiac magnetic resonance (CMR) in 56% of 400 ATTR CM patients studied.^79^ A 2018 study by Clemmensen et al. reported similar findings, with clinical cardiac amyloidosis patients demonstrating increased myocardial oxygen consumption and a reduction in myocardial external efficiency correlating with disease progression on echocardiography.^80^ Dorbala et al. observed coronary microvascular dysfunction and angina symptoms in cardiac amyloidosis patients in the absence of epicardial disease.^81^

Considering our data in light of these observations, we offer a new hypothesis for ATTR amyloid pathophysiology: dysregulated angiogenesis in the setting of thrombo-inflammatory microcirculatory occlusion. Capillary clotting, perhaps driven by endothelial cell activation via contact with unstable, misfolded TTR,^76^ creates a localized hypoxic environment that triggers angiogenesis to revascularize the hypoxic tissue. In an appropriate response, we hypothesize that stable TTR would bind to and upregulate LRP1 to facilitate organized angiogenesis and proper healing. However, if unstable WT TTR or genetic variant TTR binds to but fails to activate LRP1, disorganized angiogenesis is favored due to the competitive inhibition and lack of upregulation of the receptor. Disorganized angiogenesis prevents quenching of the hypoxic signal, perpetuating TTR migration to the area with a dearth of target receptors. Vascular permeability increases in the thromboinflammatory environment of endothelial cell activation,^82^ and the lack of vessel walls at the level of the capillary exposes the vascular basement membrane. Localized high TTR concentrations in the setting of vasodilation and increased capillary permeability lead to TTR extravasation, where misfolded TTR interacts with various components of the basement membrane, and particularly COLVI, resulting in progressive TTR conformational conversion into TTR cross-β-sheet amyloid fibrils and congestion of the vascular basement membrane. Such limitations would impede cell-to-cell communication and prevent organized revascularization, leading to the pathological remodeling and reduction of cardiac reserve observed in ATTR cardiomyopathy. Notably, closure of upstream precapillary sphincters would prevent clinical observation of microvascular occlusion in flow studies such as an angiogram. Since tafamidis substantially reduces structurally heterogeneous TTR aggregates in circulation,^41,63^ it could also substantially reduce further amyloid deposition by conformational conversion in the extracellular matrix in symptomatic CM patients.

Recent work by Levites et al.^83^ utilized unenriched proteomics in a mouse model of Alzheimer’s disease (AD) to identify proteome changes in response to amyloid β (Aβ) amyloid in the tissue. Additionally, this work incorporated proteomic analysis from Seifar et al.^84^ to provide further evidence for key signaling networks that are also present in human AD brain tissue. Our ATTR heart tissue proteomics data shows significant overlap with the proteomic profiles in both AD mice and human AD brain tissues. Critically, key amyloid interacting proteins, as well as coagulation, complement, and angiogenesis proteins described herein, were significantly changed hits (p-value <0.05) in the AD mouse, AD human brain, WT ATTR CM (vs. control), and V122I ATTR CM (vs. control) data sets. These include the amyloid signature protein APOE, angiogenesis proteins ADAMTS1, SFRP1, and VCAN, and coagulation and complement cascade protein FGG. Significantly altered in the AD mouse, AD human brain, and WT ATTR (vs. control) data sets are amyloid signature protein VTN, and coagulation and complement cascade proteins FGA and FGB. Overall, the WT ATTR and V122I ATTR data sets overlap 27% of their significant hits with both Levites et al.^83^ and Seifar et al.^84^ Individually, our data sets overlap with the Levites et al.^83^ AD mouse data by 36% and overlap with the Seifar et al.^84^ AD human brain data set by 57%. Our study utilized the IP2 analysis pipelines to identify proteins, whereas the AD studies^83,84^ used FragPipe. Reanalysis using FragPipe may yield additional hits in our data, and may explain why the key hits of interest identified in the AD data,^83,84^ e.g. MDK and PTN, are missing from our ATTR data. Thus, it seems that while transthyretin amyloidosis and AD are distinct proteinopathies, there may be a shared proteomic response to amyloidosis in tissue.

Further studies should be conducted to elucidate TTR’s role in angiogenesis and its interaction with LRP1. The relationship between TTR tetramer stability and LRP1 activation is especially important to understand, given that TTR tetramer dissociation is the rate-limiting step in ATTR aggregation and fibrillogenesis and LRP1’s association with other amyloid diseases. Such investigations will be critical in understanding the long-term safety of TTR silencer therapy as well as the potential utility of tafamidis beyond cardiac ATTR therapy.

If this hypothesis is correct, we would expect tafamidis treatment to be most beneficial early in the disease process, before vascular membrane congestion prevents appropriate revascularization by limiting the available space for sprouting angiogenesis. Further investigation into the role of stable TTR tetramer in angiogenesis could also indicate a role for tafamidis in the treatment of microangiopathies requiring revascularization.

### Limitations

This study is a clinical snapshot of end-stage sporadic and hereditary ATTR CM wherein patients exhibit advanced pathology; at the level of our study, these ATTR CM pathologies were largely indistinguishable. Further microvascular imaging studies on early-stage cardiomyopathy patients should be employed to scrutinize our hypotheses and to strengthen our understanding of the pathology cascade. While the lack of inter-patient variability information in our pathologic imaging and proteomic data renders our observations exploratory, the unenriched proteome tissue upregulation in response to TTR aggregation in the heart tissue and the proteome upregulation in response to amyloid β aggregation in human brain tissue exhibits an overlap of 57% in terms of identified proteins, including those proteins linked to coagulation, complement, and angiogenesis, justifying further scrutiny of the proposed hypotheses. Collecting additional early stage ATTR CM hearts and more later stage hearts is further motivated by the microangiopathy with thromboinflammation in the sural nerves observed in late-stage A97S TTR polyneuropathy patients.

### Conclusion

This is the first study utilizing the tissue clearing approach to image thick cardiac samples three-dimensionally to visualize the pathology of cardiac amyloidosis in humans. When paired with proteomic analysis of the visualized samples, collectively these data provide a qualitative blueprint and parts list to build a pathophysiological narrative of ATTR CM. In utilizing these methodologies in samples of both sporadic (wild-type) and genetically inherited ATTR CM (V122I), we were able to demonstrate loss of vascular architecture and regions of hypervascularization and hypovascularization present in end-stage ATTR CM. Phenotypic convergence between WT ATTR CM and V122I ATTR CM was observed by both proteomics and 3D imaging, and was further supported by amyloid fibril structural characterization by cryo-EM, as well as involvement of extracellular matrix components like COLVI (it is important to note, however, that the fibril extraction and purification process could bias our structural view of amyloid fibrils toward those fibril forms that are extracted most efficiently). Additionally, we visualized thrombosis at the level of the capillary vessels (diameter < 10 µm) in a coagulopathic environment. Further validation utilizing a larger cohort and samples in varying stages of disease is necessary to strengthen the microangiopathy hypothesis of ATTR CM disease. However, these findings have been corroborated by clinical studies with observed hypoxia and capillary loss, suggesting that the presence of hypervascularization and hypovascularization is driven by hypoxia-induced angiogenesis. While the lack of interpatient variability information in the imaging and proteomics results renders the conclusions exploratory, our imaging data and the proteomic similarity of the tissue proteome response to amyloid β and TTR aggregation has generated several hypotheses meriting further scrutiny.

## Acknowledgements

This project was funded in part by NINDS grant NS095892 awarded to Gabriel C. Lander and NIDDK grant DK046335 awarded to Jeffery W. Kelly. Jan-Hannes Schäfer received funding from the German Research Council project number 556478029. Victoria Nudell provided training and feedback for early tissue clearing experiments. The authors thank Jean-Christophe Ducom and Charles Bowman at Scripps for computational support. Emily P. Bentley provided expert editorial feedback.

