## Supplementary Information for "3D Visualization and Proteomic Analysis of Human Cardiac Transthyretin Amyloidosis Tissue Reveals Microangiopathy and Capillary Occlusion"

**SHORT TITLE:** Microangiopathy in ATTR Cardiac Amyloidosis

**Table S1.** Antibodies and dyes used in optically cleared tissue staining and western blots.

| **Antibody/Dye Target (Host species)** | **Manufacturer** | **Catalog No.** | **Dilution** | **Clone** |
|---|---|---|---|---|
| AmyTracker 480 | Ebba Biotech | A480-A-100 | 1:1000 | N/A |
| Non-native Transthyretin 5D3 (mouse) | Protego Biopharma | -- | 1:500 | Poly |
| Transthyretin Polyclonal Antibody | Invitrogen | PA-35315 | 1:2000 | Poly |
| Collagen IV (goat) | EMB Millipore Corp. | AB769 | 1:500 | Poly |
| CD144/VE-Cadherin (mouse) | Biogems | 16511-20 | 1:500 | 16B1 |
| Coagulation Factor II/Thrombin (rabbit) | Novus Biologicals | NBP1-58268 | 1:500 | Poly |
| Anti-mouse Alexa Fluor 568 (rabbit) | Invitrogen | A11061 | 1:500 | Poly |
| Anti-mouse Alexa Fluor 568 (goat) | Invitrogen | A11004 | 1:500 | Poly |
| Anti-rabbit Alexa Fluor 647 (chicken) | Invitrogen | A21443 | 1:500 | Poly |
| Anti-goat Alexa Fluor Plus 647 (donkey) | Invitrogen | A32849TR | 1:500 | Poly |

**Table S2.** Cryo-EM data collection, refinement, and modeling statistics for ex-vivo WT and V122I ATTR fibrils, COLVI microfibrils, and ATTR-COLVI co-fibrils.

| | **wt ATTR apex1** | **wt ATTR apex2** | **V122I ATTR apex** | **COLVI** | **ATTR-COLVI** |
|---|---|---|---|---|---|
| PDB ID | 9PX6 | 9PX7 | 9PX9 | NA | NA |
| EMDB ID | EMD-71953 | EMD-71960 | EMD-71962 | EMD-73313 | EMD-73312 |
| EMPIAR ID | 12909 | 12911 | 12912 | 12909 | 12909 |
| **Data Collection** | | | | | |
| Microscope | Talos Arctica | Talos Arctica | Talos Arctica | Talos Arctica | Talos Arctica |
| Camera | Falcon 4i | Falcon 4i | Falcon 4i | Falcon 4i | Falcon 4i |
| Magnification (nominal) | 150,000 | 190,000 | 190,000 | 150,000 | 150,000 |
| Voltage (keV) | 200 | 200 | 200 | 200 | 200 |
| Data acquisition software | EPU | EPU | EPU | EPU | EPU |
| Exposure navigation | image shift 8 μm | image shift 8 μm | image shift 8 μm | image shift 8 μm | image shift 8 μm |
| Electron exposure ($e^{-}Å^{-2}$) | 50 | 50 | 50 | 50 | 50 |
| Exposure rate ($e^{-}$/px/s) | 11.07 | 11.07 | 10.81 | 11.07 | 11.07 |
| Frames per movie | 1,395 | 1,395 | 1,386 | 1,395 | 1,395 |
| EER Fractions | 40 | 40 | 40 | 40 | 40 |
| Pixel size (Å/px) | 0.94 | 0.94 | 0.94 | 0.94 | 0.94 |
| Defocus range (μm) | -2.0 to -0.8 | -2.0 to -0.8 | -2.0 to -0.8 | -2.0 to -0.8 | -2.0 to -0.8 |
| Recorded movies | 3,289 | 6,232 | 6,063 | 3,289 | 3,289 |
| **Image Processing** | | | | | |
| Processing software | cryoSPARC (v.4.6) | cryoSPARC (v.4.6) | cryoSPARC (v.4.6) | cryoSPARC (v.4.7) | cryoSPARC (v.4.7) |
| Final particle images | 171,067 | 231,893 | 335,693 | 48,468 | 18,345 |
| Symmetry imposed | C1 | C1 | C1 | C1 | C1 |
| Helical twist | -1.26° | -1.28° | -1.24° | NA | NA |
| Helical rise | 4.85 | 4.85 | 4.85 | NA | NA |
| Resolution (global) (Å) | | | | | |
| FSC 0.143 (unmasked/masked) | 3.8/3.2 | 3.9/3.4 | 3.6/3.0 | 12/9.7 | 9.9/5.3 |
| Local Resolution Range | 2.7-4.9 | 3.0-5.2 | 2.6-4.8 | 7.1-12.2 | 4.2-11.9 |
| cFAR | 0.31 | 0.27 | 0.28 | 0.24 | 0.05 |
| Map sharpening B-factor ($Å^2$) | -46 | -68 | -36 | -20 | -35 |
| **Model Refinement** | | | | | |
| Initial model | PDB-8G9R | PDB-8G9R | PDB-8G9R | | |
| Refinement | Coot / ISOLDE Phenix | Coot / ISOLDE Phenix | Coot / ISOLDE Phenix | | |
| FSCmap-to-model(0.5) (Å) | 3.5 | 4.0 | 3.4 | | |
| MolProbity score | 2.04 | 2.05 | 1.91 | | |
| Clash Score | 9.80 | 8.86 | 10.01 | | |
| Composition | | | | | |
| Chains | 6 | 6 | 6 | | |
| Atoms | 8,592 | 8,592 | 8,592 | | |
| Protein residues | 546 | 546 | 546 | | |
| Ligands | - | - | - | | |
| Bonds (R.M.S.D.) | | | | | |
| Length (Å) | 0.003 | 0.002 | 0.003 | | |
| Angles (°) | 0.617 | 0.447 | 0.774 | | |
| B-factors (min/max/mean) | | | | | |
| Protein residues | 59.89/114.22/80.76 | 53.55/99.03/73.44 | 56.53/107.28/79.99 | | |
| Ligand | - | - | - | | |
| Ramachandran plot (%) | | | | | |
| Favored | 90.80 | 91.95 | 94.25 | | |
| Allowed | 9.20 | 8.05 | 5.75 | | |
| Outliers | 0.00 | 0.00 | 0.00 | | |
| Rotamer outliers (%) | 0.00 | 0.00 | 0.00 | | |
| Q-score | 0.70 | 0.67 | 0.69 | | |

**Table S3 (see Excel file).** Proteomics data of cardiac WT and V122I ATTR CM homogenates and control.

**Table S4 (see Excel file).** Proteomics data of extracted ATTR CM amyloid fibrils.

**Table S5 (see Excel file).** Complete set of tryptic, semi-tryptic, and other TTR peptides detected in fibril extracts from the WT ATTR heart.

**Table S6.** Literature Review of Evidence of TTR's Role in Angiogenesis and its Interaction with Endothelial Cells.

| Evidence of TTR's Role in Angiogenesis and Interaction with Endothelial Cells | | | |
|---|---|---|---|
| **Authors** | **Model** | **Study** | **Main Findings** |
| Obayashi K, et al.[55] | Human (in vivo) | 15 V30M FAP patients, 12 healthy volunteers | Linear negative correlation between enothelium-dependent vasodilation & disease duration; results suggest FAP peripheral vasomotor dysfunction dependent on amount of amyloid deposited around blood vessels. |
| Nunes RJ, et al.[9] | Human (in vitro) | Endothelial Cell Culture | V30M TTR decreases expression of genes involved in endothelial cell proliferation, migration, and survival compared to WT TTR |
| Lee CC, et al.[8] | Human (ex vivo), Mouse | Lung biopsy (human), Stat3C bitransgenic mice | TTR increases angiogenesis by enhancing endothelial cell tube formation and increasing endothelial cell proliferation, migration, and permeability |
| Gu Y, et al.[56] | Human (in vitro) | Endothelial Cell Culture | Formation of TTR-hnRNPA2B1 in nucleus plays role in TTR's anti-angiogenesis functions in hyperglycemia |
| Hashimoto Y, et al.[57] | Human (in vivo) | 22 cardiac WT ATTR patients, 22 one-by-one matched controls | Flow-mediated vasodilation (FMD) was significantly greater in WT ATTR patients compared to controls, with significant relationship observed between FMD and NT-proBNP in WT ATTR patients |
| Liu L, et al.[58] | Human (in vitro) | Endothelial Cell Culture | VEGFA neutralizes function of TTR's function on cell proliferation, apoptosis, migration, & angiogenesis in hyperglycemia |
| Gião T, et al.[10] | Human (in vitro), Mouse | Endothelial cell culture (human), AD transgenic mice & non-transgenic mice | TTR increased expression of pro-angiogenic molecules in cell culture; TTR reduces vascular basement membrane thickness in AD mice; TTR with stabilizer (iododiflunisal) → greater reduction of BM thickness & increased blood vessel length compared to untreated |

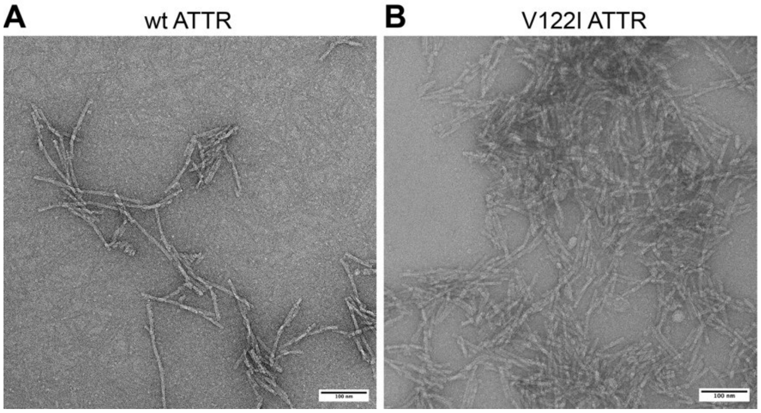


**Figure S1: Negative staining analyses of ex-vivo ATTR. (A)** Representative negative stain micrographs of wild-type (wt) ATTR. **(B)** Micrograph of ATTR amyloids of variant V122I. Scale bar corresponds to 100 nm.

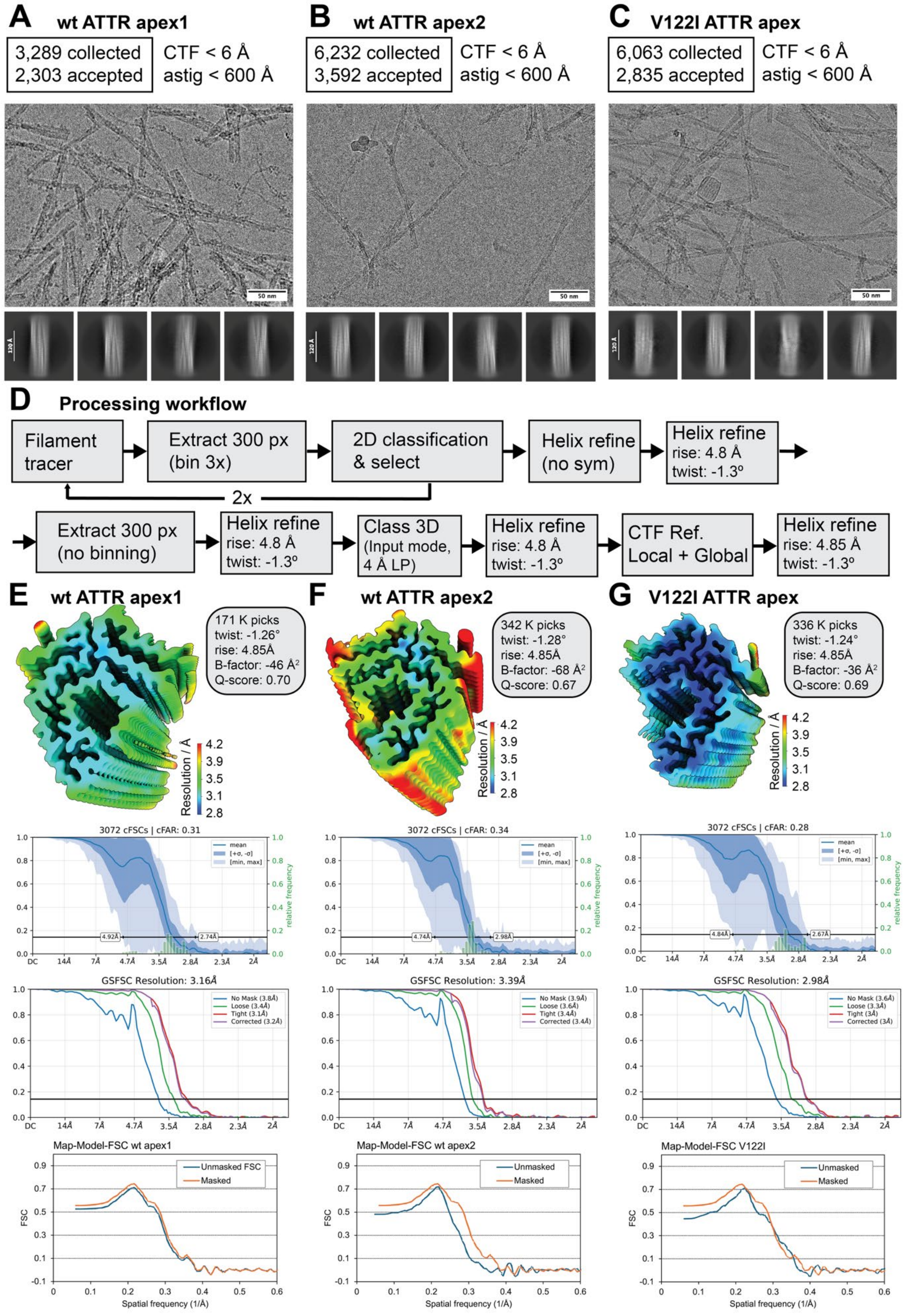
A wt ATTR apex1
3,289 collected
2,303 accepted
CTF < 6 Å
astig < 600 Å
50 nm
120 Å
B wt ATTR apex2
6,232 collected
3,592 accepted
CTF < 6 Å
astig < 600 Å
50 nm
120 Å
C V122I ATTR apex
6,063 collected
2,835 accepted
CTF < 6 Å
astig < 600 Å
50 nm
120 Å
D Processing workflow
Filament tracer
Extract 300 px (bin 3x)
2D classification & select
2x
Helix refine (no sym)
Helix refine rise: 4.8 Å twist: -1.3º
Extract 300 px (no binning)
Helix refine rise: 4.8 Å twist: -1.3º
Class 3D (Input mode, 4 Å LP)
Helix refine rise: 4.8 Å twist: -1.3º
CTF Ref. Local + Global
Helix refine rise: 4.85 Å twist: -1.3º
E wt ATTR apex1
171 K picks
twist: -1.26°
rise: 4.85Å
B-factor: -46 Å²
Q-score: 0.70
Resolution / Å
F wt ATTR apex2
342 K picks
twist: -1.28°
rise: 4.85Å
B-factor: -68 Å²
Q-score: 0.67
G V122I ATTR apex
336 K picks
twist: -1.24°
rise: 4.85Å
B-factor: -36 Å²
Q-score: 0.69
3072 cFSCs | cFAR: 0.31
3072 cFSCs | cFAR: 0.34
3072 cFSCs | cFAR: 0.28
mean
[+σ, -σ]
[min, max]
relative frequency
4.92Å
2.74Å
4.74Å
2.98Å
4.84Å
2.67Å
GSFSC Resolution: 3.16Å
No Mask (3.8Å)
Loose (3.4Å)
Tight (3.1Å)
Corrected (3.2Å)
GSFSC Resolution: 3.39Å
No Mask (3.9Å)
Loose (3.6Å)
Tight (3.4Å)
Corrected (3.4Å)
GSFSC Resolution: 2.98Å
No Mask (3.6Å)
Loose (3.3Å)
Tight (3Å)
Corrected (3Å)
Map-Model-FSC wt apex1
Unmasked FSC
Masked
Map-Model-FSC wt apex2
Unmasked
Masked
Map-Model-FSC V122I
Unmasked
Masked
FSC
Spatial frequency (1/Å)

**Figure S2: Cryo-EM data collection, image processing and validation of ATTR fibrils. (A-C)** Representative micrographs (scale bar at 50 nm) and 2D class averages (scale bar at 120 Å) with their respective data collection statistics and applied selection criteria. **(D)** cryoSPARC processing scheme used for all datasets, with additional details in sup. Table S5 and the method section. **(E-G)** Validation of the helical reconstructions, with final particle numbers, applied helical symmetry parameters, B-factor (autofit from the Guinier plot within RELION 5), and Q-score calculated in ChimeraX. The local resolution distribution was calculated in cryoSPARC using the FSC 0.5 criterion. Gold-standard Fourier-shell correlation reporting the overall resolution at FSC cutoff 0.134. Conical FSC report (cFAR) and resolution range. Map-to-Model FSC calculation from MolProbity within Phenix.

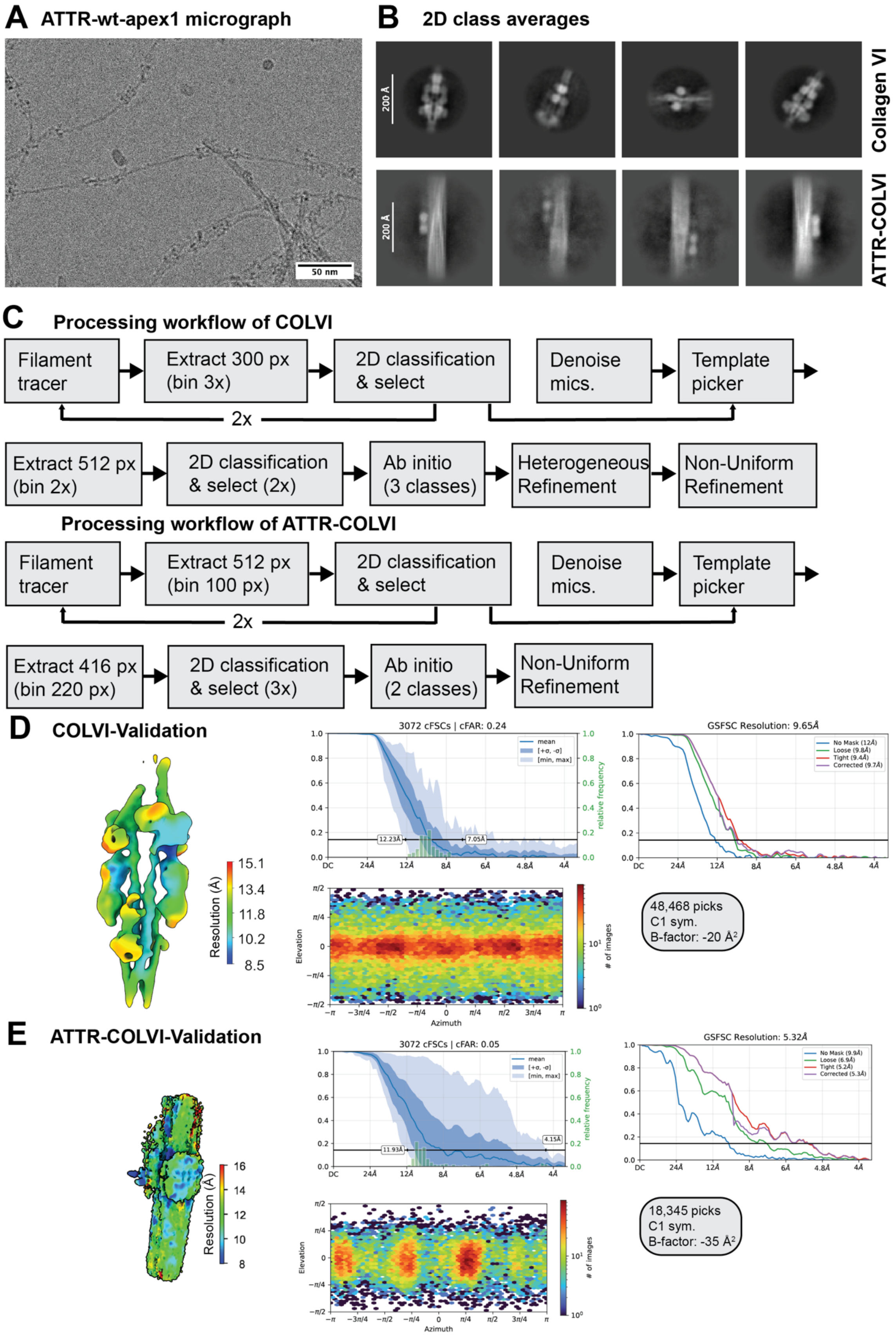
A ATTR-wt-apex1 micrograph
50 nm
B 2D class averages
200 Å
Collagen VI
ATTR-COLVI
C Processing workflow of COLVI
Filament tracer
Extract 300 px (bin 3x)
2D classification & select
2x
Denoise mics.
Template picker
Extract 512 px (bin 2x)
2D classification & select (2x)
Ab initio (3 classes)
Heterogeneous Refinement
Non-Uniform Refinement
Processing workflow of ATTR-COLVI
Filament tracer
Extract 512 px (bin 100 px)
2D classification & select
2x
Denoise mics.
Template picker
Extract 416 px (bin 220 px)
2D classification & select (3x)
Ab initio (2 classes)
Non-Uniform Refinement
D COLVI-Validation
Resolution (Å)
15.1
13.4
11.8
10.2
8.5
3072 cFSCs | cFAR: 0.24
12.23Å
7.05Å
GSFSC Resolution: 9.65Å
No Mask (12Å)
Loose (9.8Å)
Tight (9.4Å)
Corrected (9.7Å)
48,468 picks
C1 sym.
B-factor: -20 Å²
E ATTR-COLVI-Validation
Resolution (Å)
16
14
12
10
8
3072 cFSCs | cFAR: 0.05
11.93Å
4.15Å
GSFSC Resolution: 5.32Å
No Mask (9.9Å)
Loose (6.9Å)
Tight (5.2Å)
Corrected (5.3Å)
18,345 picks
C1 sym.
B-factor: -35 Å²
Elevation
Azimuth
### of images
relative frequency

61

**Figure S3. Cryo-EM data collection, image processing and validation of collagen VI (COLVI) microfibrils and ATTR-COLVI co-fibrils. (A)** Representative micrographs (scale bar = 50 nm) and **(B)** 2D class averages (scale bar at 200 Å). **(C)** cryoSPARC processing scheme used for COLVI and ATTR-COLVI, with additional details in sup. **Table S5** and the method section. **(D-E)** Validation of the reconstructions, with final particle numbers, applied symmetry parameters, and B-factor (autofit from within cryoSPARC). The local resolution distribution was calculated in cryoSPARC using the FSC 0.5 criterion. Gold-standard Fourier-shell correlation reporting the overall resolution at FSC cutoff 0.143. Conical FSC report (cFAR) and resolution range.

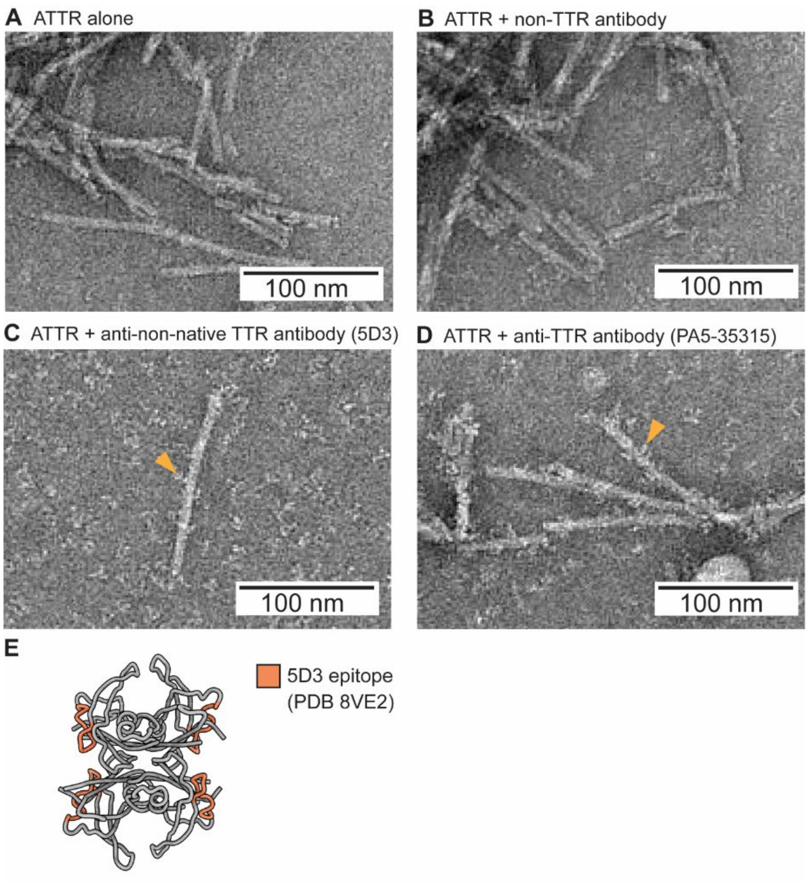


**Figure S4. Antibody binding to ATTR amyloids in negative-stain TEM.** (**A**) TEM micrograph of patient ATTR amyloid fibrils. (**B**) Negative control TEM micrograph of patient ATTR amyloid fibrils with the addition of an antibody that is not specific for TTR (anti-α-tubulin DM1A), showing that non-TTR antibodies to not recognize the ATTR fibrils. (**C**) TEM micrograph of patient ATTR amyloids in complex with the anti-non-native TTR antibody 5D3, showing that this antibody does recognize ATTR fibrils despite not recognizing native TTR. (**D**) Positive control TEM micrograph of patient ATTR amyloids in complex with a commercial anti-TTR antibody (PA5-35315, which recognizes TTR residues 71-98). Scale bar on micrographs = 100 nm. (**E**) Native soluble TTR (PDB ID 8VE2) with the 5D3 epitope mapped onto the cartoon representation of the tetramer structure (orange).

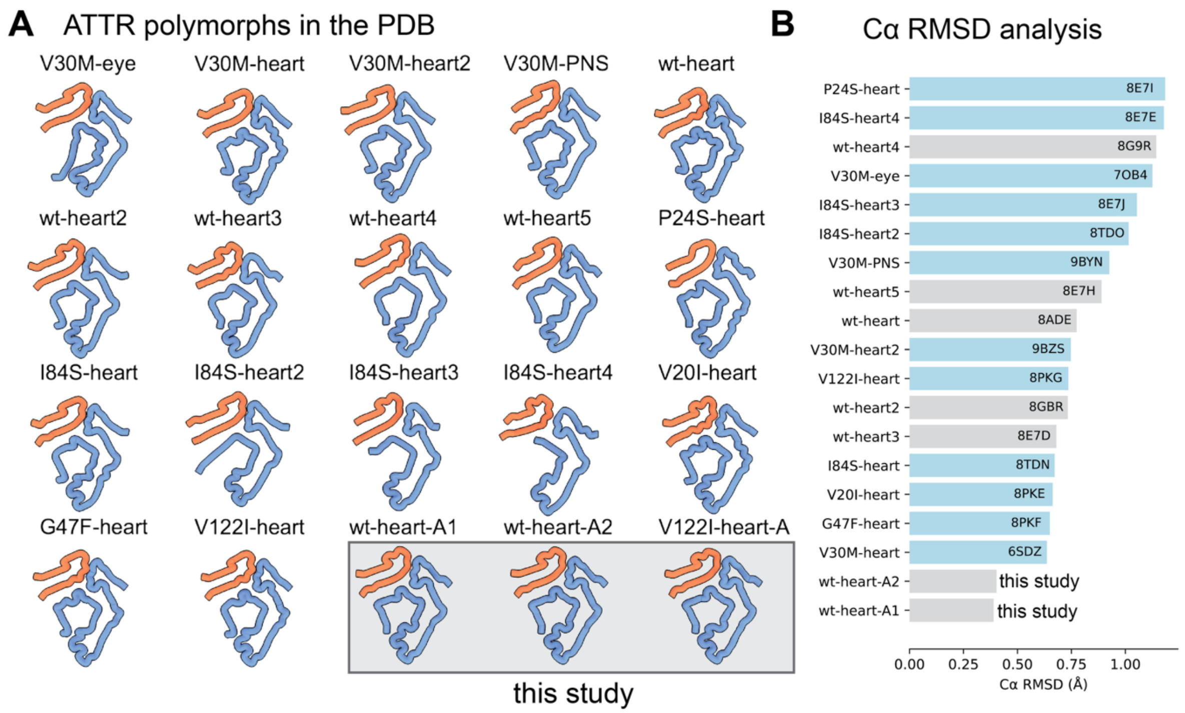


**Figure S5: Comparison of ATTR polymorphs. (A)** Z-stack models of all resolved ATTR polymorphs from the Protein Databank (PDB). For all, a single chain A was depicted, and multi-fibril assemblies were split. The smaller N-terminal peptide is shown in orange, and the larger C-terminal peptide is shown in blue. Reconstructions from this study are boxed in grey. (B) Cα root-mean square deviation between (RMSD) all chains in (A) and V122I-heart from this study, sorted by RMSD values. Wildtype (wt) reconstructions are shown in grey and familiar ATTR variants in blue. PDB IDs are shown as bar labels.

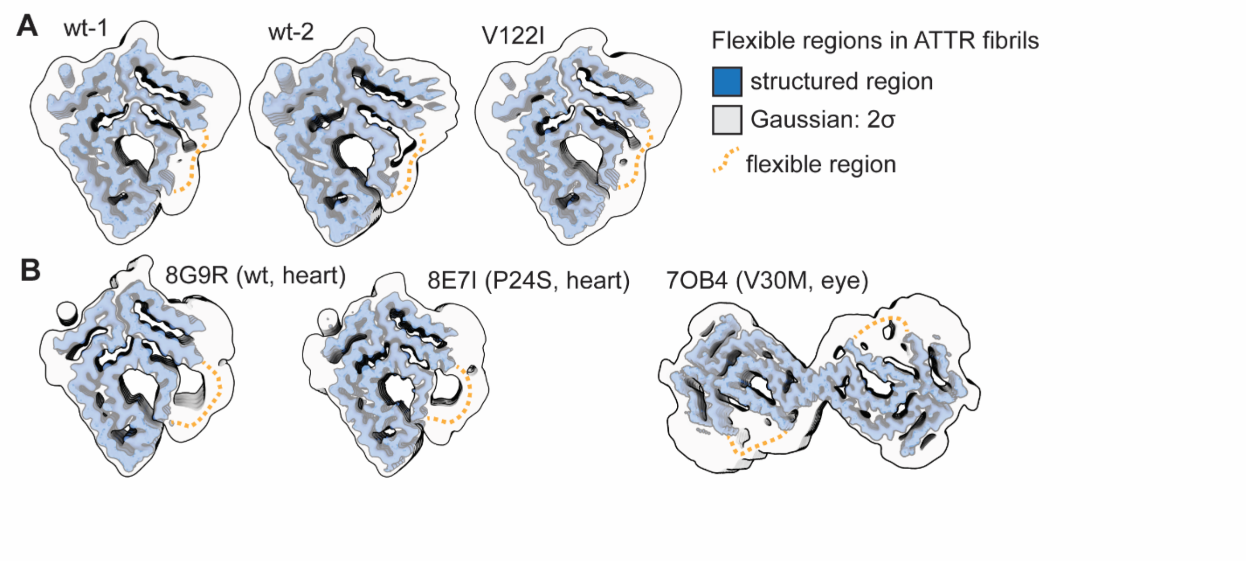


**Figure S6. Flexible regions in different ATTR fibrils. (A)** Cross-beta xy slice of the resolved cryo-EM densities for cardiac ATTR amyloids from wildtype (wt) and familiar V122I TTR variant, presented in this study. **(B)** Previously resolved wt (PDB 8G9R) and P24S (PDB 8E7I) ATTR amyloids from heart tissue and variant V30M from the vitreous body of the eye (PDB 7OB4). Structured regions, which were used for atomic-model building are shown in blue. An applied two sigma gaussian filter shows the less resolved regions within the amyloids. The flexible region spanning residues A36-G57 is marked as a dashed orange line. The 5D3 TTR antibody epitope is highlighted in blue and capitalized amino acid sequence.

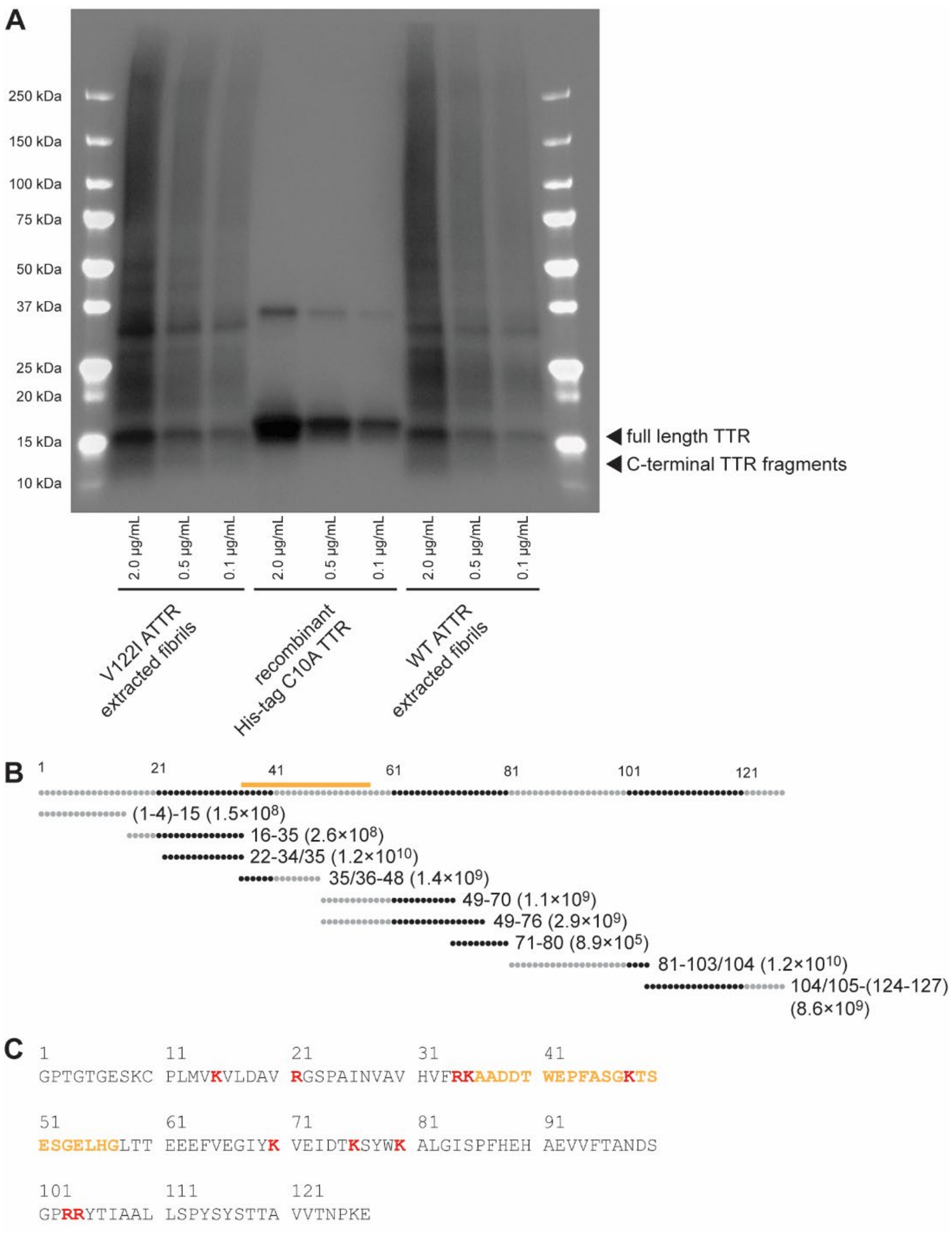


**Figure S7. ATTR fibrils from patient hearts are mostly composed of full length TTR, with proteolyzed TTR as a minor component. (A)** Western blot of extracted ATTR fibrils visualized with an antibody to the expected C-terminal fragment of TTR (residues 71-98; PA5-35315, ThermoFisher). Recombinant, His-tagged C10A TTR in the central lane is for comparison. It runs slightly heavier than the full-length TTR in the ATTR fibrils because of the His tag. **(B)** A representation of the TTR sequence showing the fragments detected by bottom-up proteomics on extracted WT ATTR fibrils. Numbers to the right of the fragments are the positions of N- and C-terminal ends of trypsin fragments (numbers in parentheses are the intensities of the detected peptides). The orange bar represents the portion of the sequence that is not observed in the cryoEM structure in Figure 6. (**C**) The sequence of TTR for reference. Tryptic cleavage sites are shown in red. The flexible portion in the cryoEM structure is shown in orange.
